# Impairment in axonal translation and cytoplasmic viscosity during aging in sensory neurons

**DOI:** 10.1101/2025.06.03.657746

**Authors:** Maria Fransiska Emily, Laurent Guillaud, Sandra De la Fuente Ruiz, Riya Agrawal, Susan Boerner, Yuto Akimoto, Tara Helmi Turkki, Marco Edoardo Rosti, Marco Terenzio

## Abstract

Mitochondria are trafficked along axons and provide the energy required for several intracellular mechanisms including molecular transport and local translation, which is believed to contribute to the homeostasis of the axonal compartment. Decline in mitochondria activity is one of the hallmarks of aging. It is still unclear, though, whether this decline corresponds to a concomitant reduction in the extent of axonal translation during aging. Using live cell imaging of sensory neurons, we found a significant decrease in the number of active mitochondria and the percentage of mitochondria localized to axons in aged mice compared to young mice. This decrease was mirrored by a loss of intracellular ATP as well as an ATP-dependent decrease in axoplasmic viscosity. In addition, the size of G3BP1 positive axonal granules and the number of FMRP axonal granules increased. Cumulatively, we found a functional decrease in the overall level of axonal translation in aged neurons. We were able to rescue this effect by increasing ATP synthesis, which induced a global decrease in axoplasmic viscosity, while promoting RNA granule solubilization and boosting axonal translation. Proteomic analysis of newly synthesized proteins in axons of aged vs young neurons revealed a dysregulation of pathways related to axonal biology and growth. We identified MAP1B and STAT3 as proteins whose axonal local synthesis was impaired in aged axons, and more notably show that this impairment could be rescued by increasing ATP synthesis. We believe that this research sheds light on axonal translation in aged neurons and its relationship with energy sources inside the axonal compartment, possibly presenting an opportunity for future therapeutics.

## Introduction

Neurons are highly polarized cells with a complex morphology, which is essential to their function. Indeed, the neurite network comprises the majority of neuronal cell volume and mediates the connections between neurons and their targets, ensuring smooth communications between the brain, spinal cord and peripheral nerves (*1*). Considering their length, which can reach meters in large vertebrates, axons cannot rely on molecular transport alone. Therefore, localized protein synthesis could constitute an important additional mechanism contributing to axonal homeostasis (*2*). Recently, an increasing body of evidence highlighted the importance of axonal translation and linked its impairment to neurodegenerative diseases (*3*). Axonally localized mRNA is bound to RNA binding proteins (RBP) in a complex often based on liquid-liquid phase separation (LLPS) (*4*, *5*). Ras GTPase-activating protein-binding protein 1 (G3BP1) and Fragile X messenger ribonucleoprotein (FMRP) have been described to form phase separated granules (*5–7*), which are known to be transported into axons, where their disassembly plays a key role in the spatial and temporal control of the translation of their associated mRNAs (*8*, *9*). Interestingly, impairment of G3BP1 granule dissolution has been linked to reduced protein translation and nerve regeneration in rats (*9*).

Mitochondria have been reported to play an active role in axonal translation, supplying the energy needed for axonal translation, as well as facilitating active transport of mRNA (*10*). Mitochondria are transported along microtubules in both directions and defects in neuronal mitochondrial health and function have been described in late onset neurodegenerative diseases prevalent with aging, such as Alzheimer’s disease, Amyotrophic Lateral Sclerosis (ALS) and Parkinson’s disease (*10*). Recently, we reported how mitochondrial impairment reduced intracellular ATP levels in human motor neurons derived from induced pluripotent stem cell (iPSC) of ALS and Parkinson patients, leading to an increase in axoplasmic viscosity and the accumulation of TAR DNA-binding protein 43 (TDP-43) aggregates in ALS axons (*11*). Interestingly, defects in mitochondrial function are also often associated with aging, as mitochondria impairment leads to reduced ATP production and increased oxidative stress (*12*). Alterations in mitochondrial axonal transport have been shown in the peripheral nervous system of aging *Drosophila melanogaster* (*13*). Subsequently, mitochondrial motility was shown to be affected with age in C. elegans motor neurons and in mouse central nervous system and peripheral neurons (*14–17*). Interestingly, oxidative stress, protein accumulation and cytoplasmic viscosity can be modulated by alteration of mitochondrial transport (*13*, *17*, *18*).

In this study, we use primary cultures of Dorsal Root Ganglia (DRG) neurons isolated from adolescent mice (8-10 weeks old) and late adulthood mice (1 year old, 52-63 weeks old) to assess changes in the extent of axonal translation in relation to aging. From now on we will address adolescent mice as young and late adulthood mice as aged. We identified a defect in the frequency and number of active axonal mitochondria in aged neurons, which was reflected in a general decrease of cytosolic ATP. This decrease in cytosolic ATP was mirrored by increased viscosity of the axonal cytoplasm of aged DRG neurons and an increase in the volume of G3BP1 and the number of FMRP positive axonal granules. We then identified a deficit in axonal translation in aged DRG neurons using a combination of microfluidics chambers and transfection of exogenous GFP mRNA in DRG axons. Importantly, treatment with NMN, an NAD^+^ precursor, was able to increase mitochondrial activity and ATP production in aged neurons, thus decreasing axoplasmic viscosity, inducing RNA granule solubilization and restoring axonal translation levels. Proteomic analysis of newly synthesized proteins in axons of aged vs young DRG neurons revealed a dysregulation of pathways related to axonal biology and growth. We chose to focus on 2 candidates, one involved in neurite outgrowth (*19*), the Microtubule Associated Protein 1B (MAP1B), and the other in the initiation of the response to axonal injury (*2*), the Signal Transducer and Activator of Transcription 3 (STAT3). The synthesis of the axonal pool of both proteins was impaired in aged DRG neurons and could be rescued by NMN treatment. Overall, our evidence points to a decrease in axonal translation in aged neurons, which is connected to the viscosity of the cytoplasm and its regulation via mitochondrial activity. This phenomenon could help to explain pathological changes observed in axons during late onset degeneration when mitochondria are affected, and suggests a possible therapeutic avenue based on metabolic regulation and modulation of cellular ATP levels.

## Materials and Methods

### Animal Experiments

All experiments involving animal subjects were carried out in accordance with the guidelines and regulations of Okinawa Institute of Science and Technology (OIST) Graduate University and approved by OIST Animal Care and Use Committee (protocol no.: 2020-304). OIST animal facilities and animal care are accredited by AAALAC International (reference no. 1551). Adult (8–10 weeks and 52-63 weeks) female ICR mice were purchased from CLEA or Jackson Laboratory. Animals were housed at 24.0 ± 0.5 °C with alternating 12 h day–night cycles and allowed access to food and water ad libitum.

### Reagents, Chemicals, and Antibodies

Culture media, sera, and chemicals were purchased from GibcoBRL, Invitrogen, and Sigma–Aldrich respectively, unless otherwise specified. Chemicals used in this study are puromycin (Sigma–Aldrich; #P8833), anisomycin (Nacalai Tesque; #03046-14), nerve growth factor (NGF) (BioVision, Inc.; #4303R-20), and β-Nicotinamide Mono-Nucleotide (Sigma-Aldrich; #N3501). Primary antibodies used in this study are α-βIII-tubulin (Synaptic Systems GmbH; #302304), α-puromycin (Merck; #MABE343), α-STAT3 (Cell Signaling Technology; #12640), α-MAP1B (Invitrogen; #PA582798), α-FMRP (Invitrogen; #PA534584), and α-G3BP1 (Sigma–Aldrich; #G6046). Secondary antibodies used for immunostaining are α-mouse Alexa Fluor 488 (Invitrogen; #A11001), α–guinea pig Alexa Fluor 568 (Invitrogen; #A11075), α–guinea pig Alexa Fluor 647 (Invitrogen; #A21450), α– guinea pig Alexa Fluor 555 (Invitrogen; #A21435), α-rabbit Alexa Fluor 488 (Invitrogen; #A11008), and α-rabbit Alexa Fluor 568 (Invitrogen; #A11011). Secondary horseradish peroxidase (HRP)–conjugated mouse and rabbit antibodies for immunoblots were α-mouse HRP-conjugated antibody (Cell Signaling; #7076) and goat α-rabbit (Abcam Limited; #ab6721).

### Microfluidics device fabrication

Microfluidics devices were constructed as described in (*20*). Briefly, PDMS (Sigma-Aldrich; #761036) was poured into a mold and baked at 60°C overnight until the PDMS was cured. The PDMS was then cut and bonded to a 35 mm glass bottom dish using a plasma cleaner in a vacuum condition under 400mTorr for 40 seconds. Once bonded, the device was sterilized using 70% ethanol for 5 minutes, washed 3 times with sterile milliQ water and exposed to UV light for 30 minutes.

### Dorsal Root Ganglia Neuronal Culture

Microfluidics chambers (MFC) or glass bottom coverslips were coated with 0.01% poly-l-lysine (Sigma–Aldrich; #P4832) overnight at 4 °C. The following day, MFCs or coverslips were washed 3 times with sterile milliQ water and further coated with laminin (GibcoBRL; #23017-015) diluted in Hanks’ balanced salt solution (HBSS) (GibcoBRL; #14175095) for 1 hour at 37 °C.

Dorsal root ganglia (DRG) neurons were dissected and disaggregated as described in (*8*). Briefly, DRGs from all segmental levels were collected in HBSS (GibcoBRL; #14175095), supplemented with 5 mM HEPES (Sigma–Aldrich; #H0887), and 0.1 mg/ml primocin (Invivogen; #ant-pm-1). Extracted DRG were then enzymatically treated first with 100 U of papain (Worthington Biochemical Corporation; #PAP) in enriched HBSS for 30 min at 37°C and then with collagenase (1 mg/ml) and dispase (1.2 mg/ml) in enriched HBSS at 37°C (Worthington Biochemical Corporation; #CLS2) (Roche Diagnostics GmbH; #04942078001) for another 30 minutes at 37°C. The ganglia were then triturated in enriched HBSS by pipetting up and down with a 1000 μl pipette. Finally, neurons were spun in a 20 % percoll gradient (Sigma–Aldrich; #P4937) in L15 medium (GibcoBRL; #L-5520), supplemented with 5 mM HEPES, 10% fetal bovine serum (FBS) (Invitrogen; #10270106), and 0.1 mg/ml primocin at 1000 rpm for 8 min, and washed twice by centrifugation in F-12 medium (Invitrogen; #11765054) supplemented with 10% FBS and 0.1 mg/ml primocin, at 1000 rpm for 3 min. Cells were plated in the MFCs or onto coverslips in F-12 medium supplemented with 10% FBS and 0.1 mg/ml primocin. When cultured in MFCs, 50 ng/ml neurite growth factor (NGF) (BioVision, Inc.; #4303R-20) was added only to the axonal side to promote axonal growth towards the axonal compartment of the MFC. After 2 days in vitro (DIV), the medium was supplemented with 2.5 μM of arabinofuranosylcytosine (Jena Bioscience GmbH; #N-20307-1) to inhibit glial proliferation.

When needed, DRG neuron cultures were transduced with AAV(PHP.eb)-CAG-GFP (SignaGen Laboratories; #SL116010) to express cytosolic GFP for live imaging of cytoplasmic viscosity and axonal morphology.

### Immunofluorescence

Neurons were fixed with 4% PFA in Phosphate Buffer Saline (PBS, GibcoBRL; #10010023) for 30 minutes and washed 3 times with PBS for 5 minutes at room temperature (RT). Cells were then permeabilized and blocked in 0.3% Triton X-100, 5% normal goat serum (Invitrogen; #10000C) in PBS for 30 minutes, incubated with primary antibodies diluted in PBS overnight at 4 °C and washed 3 times with PBS for 10 minutes at RT. Cells were subsequently incubated with secondary antibodies diluted in PBS for 1 hour at RT before being washed 3 times with PBS for 10 min at RT and mounted on glass slides using Fluoromount-G^TM^ (Invitrogen; #00495802). Primary and secondary antibody dilution in PBS were the following: α-βIII-tubulin (1:500), α-puromycin (1:2000), α-STAT3 (1:1000), α-MAP1B (1:1000), α-FMRP (1:500), α-G3BP1 (1:500), α-mouse Alexa Fluor 488 (1:500), α–guinea pig Alexa Fluor 568 (1:500), α–guinea pig Alexa Fluor 647 (1:500), α–guinea pig Alexa Fluor 555 (1:500), α-rabbit Alexa Fluor 488 (1:500), and α-rabbit Alexa Fluor 568 (1:500).

### Quantification of the number of active mitochondria in axons and of axonal diameter

DRG neurons expressing cytosolic GFP (cGFP) were transfered in Tyrode’s solution (Sigma-Aldrich; #T2397), and further incubated with TMRE (Mitochondrial membrane potential assay kit #ab287864, Abcam) for 15 minutes at 37°C before live imaging on an LSM900 confocal microscope. Volume rendering of confocal stacks was performed in Imaris 10 (Bitplane Oxford Instruments) using cGFP to delineate the axonal network and TMRE to detect mitochondria. The number, volume and activity of mitochondria were determined and compared between young and aged neurons. The estimation of axonal diameter was determined using the Imaris Filament tracer plugin.

### Measurement of intracellular ATP levels

Intracellular levels of ATP from young and old DRG neurons in culture were estimated using CellTiterGlo 2.0 kit (Promega; #G9241) and VICTOR Nivo multimode plate reader (PerkinElmer), according to the manufacturer’s instructions. Briefly, young and aged DRG neurons were cultured in 24 wells (30000 cells/wells) plates or 35 mm culture dishes (60000 cells/dish) for 2 weeks. Culture medium was then replaced with Tyrode’s solution and an equal volume of CellTiterGlo reagent was added to the well or dish. After incubation for 12 minutes at room temperature, luminescence was recorded. For comparison, the luminescence values were adjusted to the number of cells between each sample.

### Assessment of cytoplasmic viscosity in DRG neurons by fluorescence recovery after photobleaching (FRAP)

Live imaging of young and aged DRG cultures expressing cytosolic GFP was performed on the laser confocal microscope LSM900 (Carl Zeiss GmbH) equipped with an on-stage incubation chamber P-set2000 (Pecon; #133-800261) at 37°C and 5% CO_2_ and with a plan-apochromat 63x oil-immersion objective (NA = 1.4, Carl Zeiss GmbH) as previsouly reported (*11*). Briefly, acquisition settings were adjusted to maintain a 1 fps acquisition speed in the region of interest and culture medium was replaced with Tyrode’s solution before live imaging. Images were acquired every second for a total of 2-3 minutes while focus was automatically maintained throughout the acquisition period using the DefiniteFocus 2.0 system (Carl Zeiss GmbH). For FRAP experiments, the bleaching of a region of interest along the axonal shaft or in the soma was started 20 seconds after the beginning of the acquisition, and fluorescence recovery in the bleached area was monitored for the remaining 2-3 minutes. Standard FRAP analysis was applied by removing background fluorescence and corrected for any fluorescence decay from an unbleached area during the acquisition. All fluorescence recovery intensity profiles were normalized against the maximum fluorescence intensity at 20 seconds and the minimum fluorescence intensity at the end of the bleach sequence.

### Quantification of RNA granules in axons of DRG neurons cultures

DRG neurons cultures immunolabelled with β3-tubulin and G3BP1 or FMRP antibodies were imaged on the laser confocal microscope LSM900. Confocal z-stack images were 3D rendered in Imaris 10 and RNA granules (G3BP1 or FMRP) were detected and quantified (number and volume) using the Spot detection plugin with an intial spot size of 0.5 μm. The total volume of the axons present in the image was estimated from the 3D rendering of the β3-tubulin channel and the number of RNA granules per mm^3^ of axon was calculated.

### Live Imaging and Analysis of Mitochondrial Transport in DRG Neurons

DRG neurons cultured inside MFCs at DIV 14 were incubated with MitoTracker^®^ Green (100nM, Invitrogen, #M7514) for 30 minutes at 37 °C. To track retrograde transport of mitochondria, the dye was added exclusively to the cell body compartment of the MFC, while to image anterograde mitochondrial transport, MitoTracker^®^ Green was added to the axonal compartment. The dye was washed with Tyrode’s solution (Thermo Scientific Chemicals; #J67593-K2) and cells were imaged on a stage-top incubation chamber (P-set2000; Pecon; #133-800 261) of a Zeiss LSM 900 confocal microscope using a 63× oil-immersion objective (plan apochromat NA = 1.4). Temperature and CO_2_ were maintained at 37°C and 5%, respectively. Time series were recorded for 200 frames with a 4 second interval on each frame with the following parameters: image size of 256×256 pixels, 4 z-stacks ( ± 500nm stacks), and 488 nm excitation laser.

Mitochondria were imaged inside the microchannels of the MFC. Velocity distribution and displacement were tracked manually using the MtrackJ plugin in the ImageJ software. Mitochondrial displacement is defined by the cumulative distance traveled by a single carrier across frames. Moving carriers are defined by carriers that were motile for more than 3 frames. Kymograph were produced by drawing a 50 μm line along an axon and the percentage of moving carriers was established using the kymographs of each movie. Retrograde movement was defined by carriers in the microgroove that were moving towards the axonal compartment, while anterograde movement was defined by carriers in the microgroove which were moving towards the cell body compartment during the acquisition.

Total number of total and active axonal mitochondria was recorded in the axonal compartment with the same setting as the mitochondrial transport acquisition. Mitochondria were indentified using the spot tracking plugin in the IMARIS 10 software (Oxford Instruments) with a spot size detection of 0.5 μm. Total mitochondria were calculated by taking the average number of mitochondria from the first and last frame.

### Puromycin labelling

DRG neurons cultured inside MFCs were treated at DIV 14 with puromycin (Sigma– Aldrich; #P8833). Puromycin was added to the axonal chamber at a concentration of 10 μM at 37°C and 5% CO_2_ for 15 minutes. As a negative control anisomycin was utilized. Cultures were incubated with 60 μM anisomycin for 30 minutes before being incubated with puromycin (10 μM) and anisomycin (60 μM) at 37°C and 5% CO_2_ for 15 minutes. The puromycin was then washed with PBS (GibcoBRL; #10010023) and replaced with F-12 medium (Invitrogen; #11765054) supplemented with 10% FBS and 0.1 mg/ml primocin (Invivogen; #ant-pm-1). Cells were fixed with 4% PFA in PBS for 30 min at RT immediately after puromycin incubation or at 30 and 60 minutes after media replacement. After fixation, neurons were permeabilized and immunostained as previously described. Images of newly synthesized proteins were acquired using a Zeiss LSM 900 confocal microscope with a 20x objective (plan apochromat NA = 1.4) and analyzed using the software ImageJ. Puromycin fluorescence intensity was normalized by the area of the neuronal mask defined by the βIII-tubulin staining, a neuronal specific marker in the context of adult DRG neurons.

### GFP mRNA *in vitro* synthesis

All recombinant DNA experiments were carried out in accordance with the guidelines and regulations of OIST genetic manipulation procedures and approved by the Biosafety Committee (protocol number: RDE-2024-050-2 & RDE-2024-048). eGFP cloned in the pcDNA3.1 plasmid (Addgene) was linearized using SmaI (New England Biolabs, #R0141S) and purified using a PCR Purification Kit (QIAGEN, #28106) according to the manufacturer’s instructions. *In vitro* transcription was performed using MEGAscript ®-T7 kit (Life Technologies, #AM1333) according to manufacturer’s instructions. Briefly, 1mg of purified linear DNA was used in a 20 μl reaction. 3’-O-Mem7G(5’)ppp(5’)G RNA Cap Structure Analog (New England BioLabs; #S1411L) at 4:1 ratio to GTP was added to the transcription mix to maximize the number of capped transcripts and ensure the stability of the mRNA during translation. The reactions were incubated for 4 hours at 37°C and further treated with 1 μl TURBO DNase^TM^ (Life Technologies; #AM2238) for 15 minutes at 37°C before purification using MEGAclear™ Transcription Clean-Up Kit (Life Technologies; #AM1908) according to the manufacturer’s instructions. The purified mRNA product was treated with Antarctic Phosphatase (New England Biolabs; # M0289S) to prevent the transcript from self-ligating and was purified one more time using MEGAclear™ Transcription Clean-Up Kit (Life Technologies; # AM1908).

### Live Imaging of Protein Translation

GFP mRNA was transfected into the axonal compartment of DRG neurons cultured inside MFCs at DIV 14 using Lipofectamine MessengerMAX (ThermoFisher Scientific; # LMRNA003) in Opti-MEM Reduced Serum Medium (Gibco, #31985062) according to the manufacturer’s instructions. Negative control groups were incubated with 40 μM of anisomycin (Nacalai Tesque; #03046–14), a protein synthesis inhibitor, for the length of the experiment. The cells where then transferred to a stage-top incubation chamber (P-set2000; Pecon; # 133-800 261) mounted on a Zeiss LSM 900 confocal microscope and imaged with a 63× oil-immersion objective (plan apochromat NA = 1.4). Time series were acquired every 30 minutes for 16 hours with the following parameters: image size of 512×973 pixels, 2 tiles per image, 20 z-stacks (1.06 μm stacks), and 488 nm excitation laser. Temperature and CO_2_ were maintained at 37°C and 5% respectively for the duration of the acquisition.The fluorescent intensity of the translational GFP foci was analyzed using a Python script developed in house. The script recognizes the fluorescence intensity of the spots and deducts the background intensity to the total fluorescence intensity of each spot. The resulting intensity value is further normalized to the total axonal area. The number of spots was determined using the IMARIS 10 imaging software. Spots were identified using the spot tracking plugin in IMARIS 10 with an initial spot size detection of 0.75 μm and automatic background subtraction, and the spot region growth was based on their absolute intensity. The number of spots was also normalized by the area of the corresponding axonal network, which was manually tracked using the ImageJ software. For visualization purposes, the axonal contour was tracked using the filament analysis plugin on IMARIS 10.

### Experimental Design and Statistical Rationale for MS

The proteomics experiment was designed to identify the translatome of cell bodies and axons of neurons from young and aged mice using protein tagging with O-propargyl-puromycin (OPP). As a negative control, neurons grown in the transwell system were treated with 60 μM anisomycin for 30 minutes prior to OPP incubation to ensure the proteins detected were translated proteins and not unspecific binding to strepatividin beads. Three biological replicates from 3 different mice were utilized to ensure statistical power of detection by variance. Three technical replicates were acquired by running each biological replicate three times to ensure optimal collection of peptide information and to reduce noise in the statistical analysis.

### Sample preparation for Mass Spectrometry Analysis

DRG neurons grown on a trans-well system at DIV 6 were tagged with OPP as described in (*21*). As a negative control, 60 μM anisomycin was added to the trans-well and incubated at 37 °C for 30 minutes prior to OPP treatment. Protein tagging was done using 10 μM OPP for 1 hour at 37 °C. Cells were then washed 3 times in PBS on both sides of the transwell. Cell body and axon were lysed and scraped separtely using RIPA lysis buffer (100 mM Tris-HCl pH 8.0, 1 mM EDTA, 1%Triton X-100, 0.1% Sodium Deoxycholate, 0.1% SDS, 140 mM NaCl in dH_2_0) supplemented with proteinase inhibitor cocktail (Roche Diagnostics GmbH; # 4693132001). Lysates were then kept on ice for 30 minutes before centrifugation at 15,000 rpm for 20 minutes at 4°C. The supernatant was kept and subjected to CLICK reaction as follows: 10% SDS (Invitrogen; #AM9820), 5 mM Biotin-PEG3-azide (Funakoshi; #23419), 2 mM TBTA (Sigma; #678937), 50 mM CuSO_4_ (Jena Bioscience; #CLK-MI004-50), and 50 mM TCEP (Sigma-Aldrich; #C4706) in a rotator for 2 hours at RT. Afterwards, proteins were precipitated with 5 volumes of ice-cold acetone (Sigma-Aldrich; #0104605) and kept at −20°C overnight. Acetone was removed by centrifugation at 3000 rpm for 15 minutes at 4°C and proteins were resuspended in 1 ml of methanol (Sigma-Aldrich; #1924205). Protein samples were sonicated for 10 minutes. Ice was added to the sonicator to keep the environment cool and prevent protein degradation. Methanol was removed by centrifugation at 3000 rpm for 15 minutes at 4°C and the pellete was air dried before resuspension with 1% SDS in PBS. After a second sonication using the same condition as previously mentioned, the samples were further centifuged at 3,000 rpm at RT to remove residual SDS. The supernatant was transfered to a low protein binding tube and 30 μl of the solution was removed for westren blot. 20 μl of strepatividin beads were prepared by two washes with 1% NP40 (Sigma-Aldrich; #NP40S) in PBS supplemented with proteinase inhibitor. The resuspended beads were added to the protein solution and incubated overnight at 4°C. The next day, the beads were immobilized and washed twice with 1% NP40 and 0.1% SDS in PBS for 10 minutes at RT. Afterwards, the beads were immobilized and washed 3 times with 6 M urea (Nacalai Tesque; #35904-45) in PBS using a rotator for 10 minutes at 4°C and transferred to new low protein binding tubes. The urea washes were followed by another 3 washes in ice-cold PBS for 10 minutes in a rotator at RT. Beads were then rinsed with 100 μl digestion buffer containing 45 mM Tris-HCL (pH 8) with 2 mM CaCl_2_. Aftewards, digestion buffer supplemented with 5 mM DTT (Wako; #042-29222) was added to the beads. The beads were incubated at 56°C for 30 minutes on a shaking block at 800-1000 rpm and let to cool to RT. 20 mM Iadoacetamide (IAA, Wako; #042-05591) was then added to the beads for alkykation and the samples were kept at RT in the dark for 15 minutes. Samples were incubated again with 5 mM DTT solution for 5 minutes at RT, followed by trypsin digestion using 500 ng of sequencing grade trypsin (Agilent; #204310-51) on a shaking heater block at 800-1000 rpm at 37°C overnight. The next morning, another 250 ng of trypsin was added and digestion was continued for another 4 hours before the reaction was quenched using 1% formic acid (Kanto Chemical; #16245-63). Digested peptides were purified with a monospin column C18 (GL Sciences Inc.; #501021700) and eluted with 95% ACN (Kanto Chemical; #01033-79) and 5% formic acid in water. The elution was dried using SpeedVac and resuspended in 10 μl of 0.1% formic acid for mass spectrometry analysis

### Mass Spectrometry acquisition and data analysis

For LC–MS/MS, peptides were analyzed on a Fusion Lumos (Thermo Fisher Scientific) in data-dependent acquisition. Peptides were separated by a nanoflow liquid chromatography system (M-Class, Waters) on a trap column (2 cm x 180 µm nanoEase M/Z Symmetry C18 Trap Column) and a separation column (15 cm x 75 µm nanoEase M/Z HSS T3 Column) at a 500 nl/min with a multistep gradient. Mobile phase A: water with 0.1% formic acid; mobile phase B: acetonitrile with 0.1% formic acid. Cleaned peptides were quantified using BCA Protein Assay Kit (Thermo Scientific; #23227) and diluted to a concentration of 100 ng/μl using 0.1% formic acid. A total of 200 ng of digested peptides were injected per triplicate in 3 separate measurements for technical replicates. Biological replicates were measured on separate days and normalized post-acquisition using R software. In brief, the 180-minute data-dependent acquisition method and LC conditions used to measure the samples were as follows: the peptides were eluted from the analytical column with a linear gradient from 5% B to 30% B. The mass spectrometer was set to perform data acquisition in the positive ion mode in the scan range MS at 380-1,500 m/z and auto for MS/MS. The cycle time was set to 3 sec, with charge states 2 to 7 selected from each survey scan and subjected to HCD fragmentation. For the MS and MS/MS scan parameter settings are as follows: collision energy, 30%; electrospray voltage, 2.0 kV; capillary temperature, 275℃.

All MS and MS/MS data were analyzed by Proteome Discoverer v3.1 (Thermo Fisher Scientific) for protein and peptide identification with Chymerys algorithms using label-free quantification methods. The data were queried against a Uniprot/SWISS-Prot database for *Mus musculus* database (UP000000589.fasta, 54,727 proteins, April 2024). All database searches were performed using a precursor mass tolerance of ± 10 ppm, fragment ion mass tolerance of ± 0.6 Da, enzyme specificity was set to trypsin, and maximum missed cleavages values of 2. Cysteine carbamidomethylation was set as fixed modification. Percolator was used to adjust the false discovery rate (FDR) to 1% on peptide level. Protein abundances were derived from precursor ion intensities using the LFQ quantification node. The raw abundance values were subjected to total peptide amount normalization, where the sum of peptide intensities in each sample was scaled to correct for sample loading differences and technical variability ensuring comparability across samples by equalizing the total signal intensity. The resulting values—reported as “Abundance (Scale)”—reflect normalized protein abundances were filtered and analysed using the R software. Two filtering steps were performed. During the first filtering step, proteins that were missing values in 2 or more technical repeats were omitted from the analysis. On the second filtering step, filtered protein abundances were normalized using median normalization and analyzed using the DeqMS plugin in R (*22*); proteins whose abundance was not significant (spectra count p-value> 0.1) against their anisomycin negative control counterparts were ommited from the analysis. The output data passing both filtering steps was normalized by dividing the abundance of each protein by the average abundance of each biological repeat. To identify abundance differences between proteins from young and aged neurons, the normalized data set was analyzed using DeqMS plugin in R. Proteins were considered upregulated or downregulated in the young and aged neurons if they achieved a spectra count p-value of < 0.05. Volcano plot and heatmap were visualized in R using EnchancedVolcano (https://bioconductor.org/packages/devel/bioc/vignettes/EnhancedVolcano/inst/doc/EnhancedVolcano.html) and pheatmap plugin (https://github.com/raivokolde/pheatmap) respectively.

### NMN Chronic Treatment

DRG neuron culture from young and aged mice at DIV 7 were treated with 1mM of β-NMN (Sigma-Aldrich; #N3501) for 6 consecutive days before image acquisition and analysis as mentioned in the immunofluorescence, proximity ligation and RNA scope sections. For neuronal culture grown on MFCs, β-NMN was added only to the axonal compartment.

### Proximity Ligation Assay

Proximity ligation assay (PLA) was performed using Duolink PLA reagents (Sigma– Aldrich) according to the manufacturer’s instructions. In the case of β-NMN treatment analysis, neurons were treated with β-NMN as described in the chronic β-NMN treatment section. DRG neurons cultured on coverslips were fixed at day in vitro (DIV) 6 and DIV 14 in 4% PFA diluted in PBS and permeabilized as mentioned in the immunofluorescence section. α-STAT3 (cell signaling technology; #12640), α-MAP1B (Invitrogen; # PA582798) and α-puromycin (Sigma–Aldrich; #P8833) primary antibodies were used for the PLA experiment. α-mouse minus and α-rabbit plus probes (Sigma–Aldrich; #DUO92004 and #DUO92002), were used to bind to the primary antibodies, and the signal was detected using the far-red detection kit (Sigma–Aldrich; #DUO92013). Cells were counterstained with α-βIII-tubulin (Synaptic Systems GmbH; #302304) for 1 hour at RT. Coverslips were mounted with Fluoromount-G^TM^ (Invitrogen; #00495802) overnight. Images were taken using a Zeiss LSM 900 confocal microscope with a 63X oil-immersion objective (plan apochromat NA = 1.4). PLA signal in the axon was quantified using ImageJ with manual thresholding with the same values for young and aged neurons. The axonal network was defined by the βIII-tubulin mask area. The number of puncta was calculated and divided by the neuronal network area after subtracting any areas of glia or debris. PLA signal in the cell body was quantified using IMARIS 10. The cell body was masked by the βIII-tubulin signal to separate PLA signal from the cell body from the axonal ones. PLA signal was detected using the spot tracking plugin in IMARIS 10 with the same spot area value and thresholding range between neurons from young and aged mice. The calculated number of puncta was divided by the masked cell body area.

### Puro-PLA

Detection of newly synthesized MAP1B and STAT3 was performed by incubating neuronal cultures with 10 μM puromycin (Sigma–Aldrich; #P8833) in F-12 for 15 min at 37 °C in a 5% CO_2_ incubator. Protein synthesis inhibition control groups were preincubated with 60 μM of anisomycin (Nacalai Tesque; #03046–14) in F-12 for 30 min before the addition of puromycin. The incubation was terminated by two quick washes in PBS, and cells were fixed immediately using 4% PFA for 30 min. After permeabilization and blocking, PLA was performed as described using α-puromycin (Merck; #MABE343) and α-MAP1B (Invitrogen; #PA582798), as well as α-STAT3 (Cell Signaling Technology; #12640). Puro-PLA signal was quantified as previously described in the PLA section.

### RNAscope on DRG neurons cultures

Neurons grown on the coverslips were transduced with AAV(PHP.eB)-CAG-GFP at DIV 2 and DIV 7. These cultures were subsequently utilized for RNAscope at DIV 6 and DIV 14, respectively. The detection workflow was performed according to the manufacturer’s instructions with minor modification. Briefly, DRG neurons were fixed in 4% PFA in PBS for 30 minutes at RT and subsequently washed with PBS 3 times. The samples were then dehydrated sequentially with 50%, 70% and 100% ethanol for 1 minute and kept in 100% ethanol at −20°C until the assay was performed. The cells were then rehydrated by submerging them in 70% and 50% ethanol sequentially for 1 minute before PBS wash. The samples were permeabilized in PBS-Tween (0.1%) for 10 minutes and treated with hydrogen peroxide (Advanced Cell Diagnostics; #22381) for another 10 minutes at RT. After PBS washes, the cells were incubated with primary antibodies (α-FMRP (Invitrogen; #PA534584), and a α-G3BP (Sigma–Aldrich; #G6046)) overnight at 4°C. The next day, cells were washed with PBS and postfixed with 4% PFA in PBS for 30 minutes at RT. After post fixation, cells were further washed with PBS and treated with protease III (Advanced Cell Diagnostics; #322381, 1:100 dilution) for 10 minutes at RT before being washed in PBS again. RNAscope was performed using multiplex v2 fluorescent *in situ* hybridization (Advanced Cell Diagnostics; #323110) as instructed by the manufacturer. Cells were hybridized with the probes (MAP1B probe; Advanced Cell Diagnostics; #1045181-C3, or STAT3 probe; Advanced Cell Diagnostics; #425641-C2), along with the postive (Advanced Cell Diagnostics; #320881) and negative control probes (Advanced Cell Diagnostics; #320871) at 40°C for 2 hours. Amp1, amp2 and amp3 hybridization cycles were performed before developing the C3-HRP channel for MAP1B probe or C2-HRP channel for STAT3 probe using TSA vivid fluorophore 650 for 30 minutes at 40°C. The last HRP blocking step was performed for 30 minutes at 40°C before the cells were incubated with the secondary antibodies for 30 minutes at RT. Coverslips were then washed with PBS and mounted with Fluoromount-G^TM^ (Invitrogen: #00495802) and imaged using a Zeiss LSM 900 confocal with Airyscan 2.0 super-resolution module using a 63× oil-immersion objective (plan apochromat NA = 1.4). Images were quantified using IMARIS 10 imaging software. Axons were identified using IMARIS surface rendering plugin on the basis of the GFP channel. G3BP1 and FMRP granules were identified using IMARIS surface rendering plugin with smooth filter and seed point diameter of 0.5 μm. A mask was created for the G3BP1 and FMRP granules were created to identify STAT3 and MAP1B mRNA that were located inside the granules. STAT3 and MAP1B mRNA were indentified using IMARIS surface rendering plugin with smooth filter and seed point diameter of 0.5 μm.

### Statistical Analysis

All experiments were performed in at least three biological replicates. Analysis of multiple groups was performed using one-way or two-way ANOVA with Tukey multiple comparison correction post-test. For two-group analyses, unpaired Student’s t-test was used. All statistical analyses were performed using Prism 10 (Graph-pad Software Inc.). Data are presented as mean ± SEM. Statistical significance tests and the number of samples used are described in the figure legends. Significance values are indicated as ∗p <0.05, ∗∗p <0.01, ∗∗∗p <0.001, ∗∗∗∗p <0.0001; n.s. indicates not significant.

### Image and Figure preparation

All confocal images were acquired with Zen blue software version 3 (Zeiss) and images were exported in TIF image format. 3D rendering was performed using IMARIS 10 (Bitplane Oxford Instruments), and the 3D images were exported in TIF format. All TIF files were further processed in the software Inkscape (gitlab.com/inkscape/inkscape) to create the final figures.

## Results

### Axonal mitochondria of DRG neurons isolated from aged mice display a trafficking and activity deficit

Mitochondrial disfunction is a well-known hallmark of aging and was recently described in primary Dorsal Root Ganglia (DRG) sensory neurons dissected from 2 year old mice (*17*). Indeed, DRG sensory neurons, unlike other neuronal types, are a good neuronal model for aging due to their ability to be cultured from adult animals. We looked at aged mice (1 year old, 52-63 weeks old) and compared them to young mice (8-10 weeks old) to confirm whether mitochondrial disfunction could be detected at this early stage of aging in cultured DRG neurons. We specifically wanted to focus on the axonal compartment to assess the overall health of the neuronal network. To fluidically isolate DRG cell bodies from their respective axonal projections, we took advantage of a microfluidics setup we developed to culture adult DRG neurons (*20*). Mitochondria were labelled using MitoTracker^®^ green to monitor their axonal retrograde and anterograde trafficking. Similarly to (*17*), we found no significant difference in the total number of axonal mitochondria (**Figure S1A**), their anterograde and retrograde speed (**Figure S1B & C** and **Figure S1B & D**) and retrograde displacement (**Figure S1E**). However, we observed a significant decrease in the anterograde displacement and percentage of motile mitochondria in DRG neurons isolated from aged mice (**Figure S1E**). In addition, while the number of total axonal mitochondria remained the same, we observed a significant decrease in axonal active mitochondria stained with TMRE in DRG neurons from aged mice compared to those isolated from young mice (**Figure 1A, B, C & D**). Taken together our observations suggest that a decline in mitochondrial activity can already be observed in the early stage of aging.

**Figure 1.**
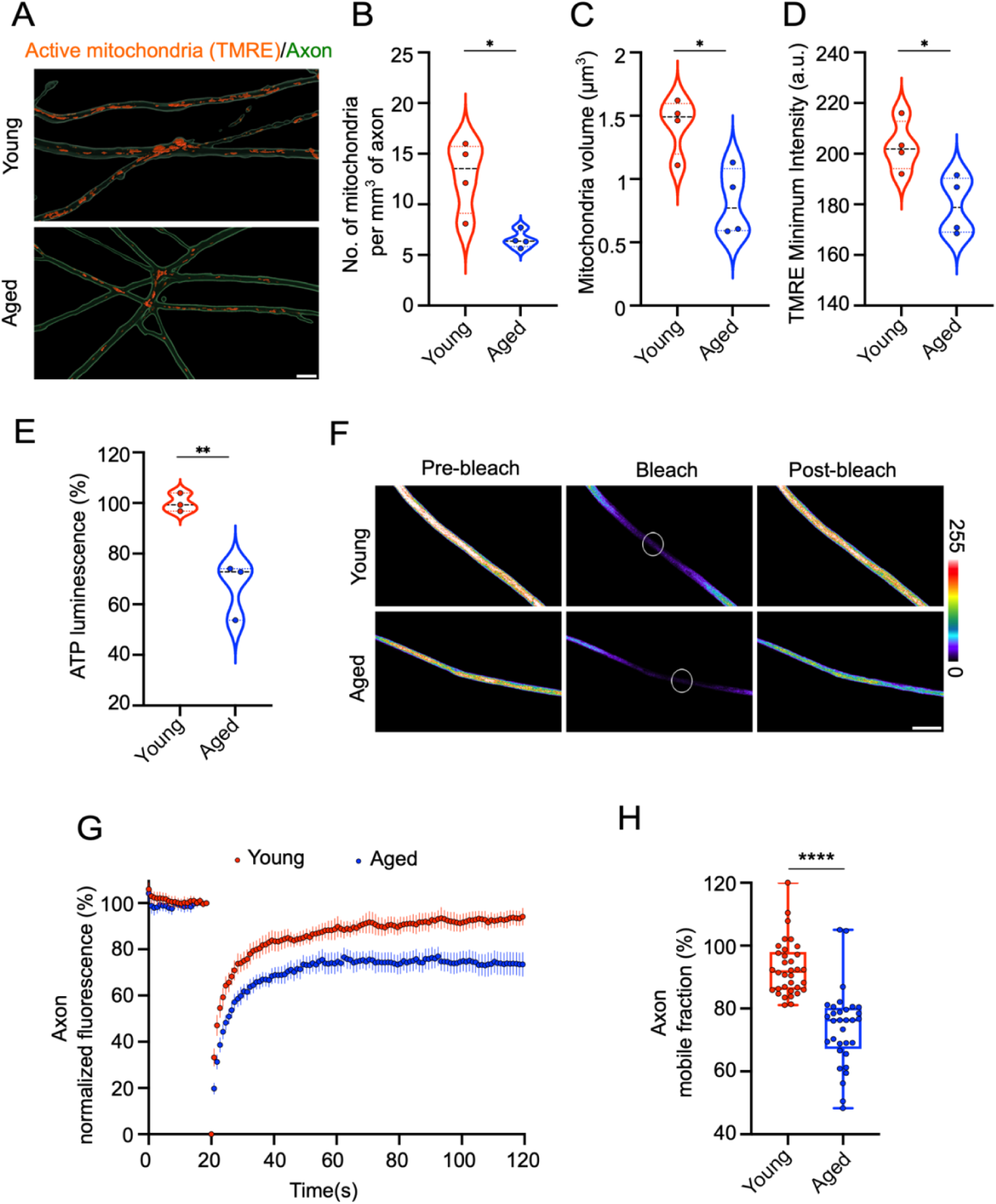
Mitochondrial and viscosity impairment in aged neurons. (A) Representative 3D rendered images of active mitochondria labeled with TMRE (red) in axon (green) of DRG neurons expressing cGFP isolated from young and aged mice. Scale bar = 5 µm. (B-D) Quantification of the number (B), volume (C) and TMRE intensity (D) of active axonal mitochondria for the experiment described in (A) (n= 4; means ± SEM; *p < 0.05 ; unpaired t test). (E) *In vitro* bioluminescence measurement of intracelllular ATP level in DRG neurons from young and aged mice (n= 3; means ± SEM; **p < 0.01 ; unpaired t test). (F) Axon of DRG neurons at DIV 14 from young and aged mice expressing cGFP (pseudo-color). Panels from left to right show ROI before photo-bleaching, during bleaching, and post recovery. Scale bar = 5 µm. The color bar represents cGFP fluorescence intensity. (G) Axonal fluorescence intensity recovery profile of DRG neurons at DIV 14 from young and aged mice. Mean ± 95% CI. (H) Mobile cGFP fraction representing cytosolic fluidity estimation from the last 20 seconds of the experiment described in (G) (young n=34, aged n= 33 from 4 biological repeats; means ± SEM; *p < 0.05 ; unpaired t test).

### Decrease in Cellular Viscosity due to ATP deficit in aged mice

In a recent work we correlated a defect in mitochondria activity with a decrease in axonal ATP leading to an increase in cytoplasmic viscosity and protein aggregation in neurons derived from human iPSC of Parkinson and ALS patients (*11*). Based on this work, we wondered whether the mitochondrial defect we observed in axons of aged DRG neurons correlates with a decrease in intracellular ATP levels as well as cytosolic fluidity. Indeed, we observed a lower concentration of intracellular ATP in DRG neurons cultured from aged mice compared to young ones (**Figure 1E**). To measure the axoplasmic fluidity of the axonal compartment we decided to use fluorescence recovery after photobleaching (FRAP) of cytosolic GFP (cGFP). Indeed, FRAP is widely used to study LLPS both *in vitro* and *in vivo* (*23*, *24*) and has been utilized to assess cytosol viscoadaptation upon energy deprivation both in yeast (*25*, *26*) and mammalian neurons (*11*). The percentage of fluorescence recovery, which correlates with cGFP’s diffusion rate, allows for an estimate of cytosolic fluidity. In this context, an higher recovery rate indicates better cytosolic solubility, while a lower one suggests a less soluble cytoplasm. FRAP analysis also revealed a lower cGFP recovery level in aged mice compared to young mice after photobleaching (**Figure 1F, G & H**). This suggests that cellular viscosity is increased with age in axons of peripheral neurons. We also observed a smaller, but significant increase in the cytosolic viscosity of the cell body of aged neurons (**Figure S2A, B & C**) in agreement with previous findings (*17*).

### Axons of DRGs from aged mice display a larger size and number of G3BP1 and FMRP positive granules, respectively

Recently, LLPS has been associated with the formation of RNA granules (*27*). These granules, characterized by the presence of RNA binding proteins such as G3BP1 or FMRP (*6*, *7*, *28*), are critical for the localization of mRNA to axons and their subsequent translation (*9*, *29*). We have previously associated an increase in axonal cytoplasmic viscosity with an increased number of TDP-43 aggregates in ALS motor neurons derived from human iPSCs (*11*). Given our findings on decreased ATP levels and increased axonal cytoplasmic viscosity in DRG neurons from aged mice (**Figure 1E-H**), we wondered whether mRNA granules could also be affected. Thus, we checked for the presence of G3BP1 positive granules in axons of DRG neurons from aged mice, as G3BP1 has been shown to be an essential component of mRNA granules (*9*). We have observed that G3BP1 positive granules in axons of DRG neurons from aged mice exhibit a larger volume compared to those from young mice (**Figure 2A & B**). In addition, our data showed that FMRP granules were similarly impacted by the increased cytoplasmic viscosity. Indeed, we detected a higher number of FMRP-positive granules in axons of aged mice compared to those from young mice (**Figure 2C & D**).

**Figure 2.**
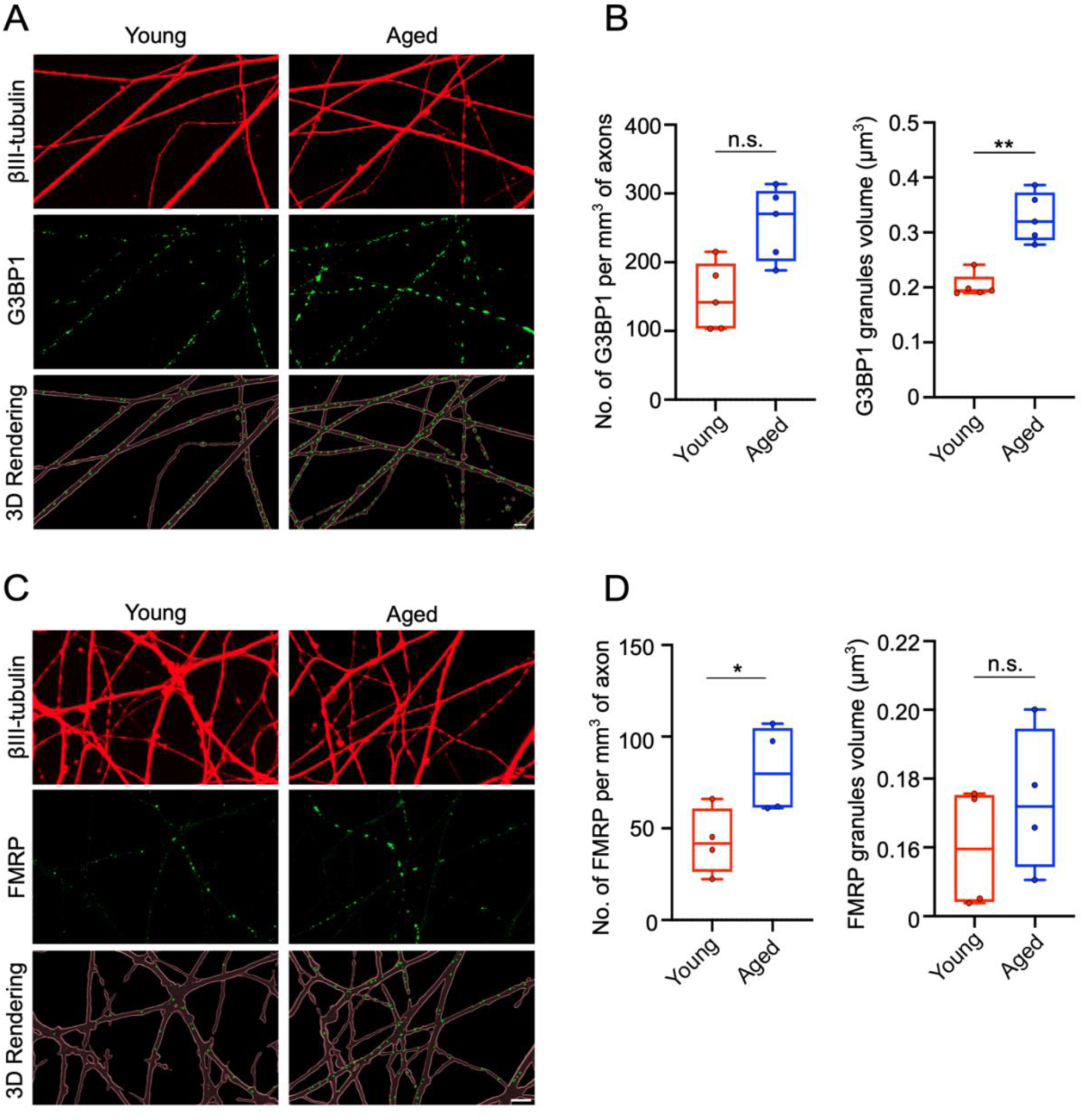
Aggregation of G3BP1 and FMRP granules in aged neurons. (A) Representative images of G3BP1 granule (in green) in axons of young and aged mice. Axons were labelled with βIII-tubulin (in red). Bottom panels represent merged 3D images of RNA granules along the axons. Scale bar = 5 μm. (B) Quantification of the number and volume of G3BP1-positive granules in DRG neurons from young and aged mice (n= 5; means ± SEM; ns, not significant, *p < 0.05 ; unpaired t test). (C) Representative images of FMRP granules (in green) in axons of young and aged mice. Axons were labelled with βIII-tubulin (in red). Bottom panels represent merged 3D images of RNA granules along the axons. Scale bar = 5 μm. (D) Quantification of number and volume of FMRP-positive granule of G3BP1-positive granule in DRG neurons from young and aged mice (n young= 34, n aged= 33; means ± SEM; ns, not significant, ****p < 0.0001 ; unpaired t test).

### Axons of DRGs from aged mice display lower levels of axonal protein translation

It has been reported that aging reduces axonal translation in axons of neurons of the central nervous system (CNS) derived from human embryonic stem cells (hESC), impacting their regeneration capacity (*30*). Thus, we tested the extent of local protein production in axons of primary sensory neurons taken from aged mice and compared it to those of axons from young mice. As previously described, we took advantage of a microfluidics setup to fluidically isolate DRG cell bodies from their respective axonal projections. We monitored protein translation using puromycin, a t-RNA analog widely used for monitoring protein translation due to its ability to incorporate into nascent polypeptide chains (*31*). We performed a pulse-chase experiment by incubating the axonal compartment with puromycin for 30 minutes, washing the cells and then fixing them at 0, 30, and 60 minutes after washing. Puromycin signal was initially detectable by immunostaining only in the axonal compartment and gradually appeared in the cell body compartment as expected (**Figure S3A & B**). Puromycin labelling of neurons cultured on MFC showed a 10% decrease in the level of axonal translation in the aged mice (**Figure S3C & D**).

One of the drawbacks of using puromycin labelling is that the cells need to be fixed to visualize the puromycin signal. To visualize axonal translation live, we synthesized *in vitro* and subsequently transfected the axons of DRG neurons with mRNA for EGFP in microfluidic devices. This method has also been previously adopted to look at translation in *Xenopus* oocyte *in vitro* (*32*). The use of the microfluidic devices enabled us to add the transfection mixture directly to the axonal compartment, excluding neuronal cell bodies, thus allowing for the visualization of axonal EGFP translation, without the confounding aspect of anterograde transport of EGFP synthesized in the soma. We then analyzed the spots using a script that was developed in house to detect the foci of translation and measure their intensity and area (**Figure S4A & B**). Using this system, we monitored the increase in EGFP fluorescence over a time course of 16 hours. When quantifying the number of translational hotspots in axons of DRG neurons from aged mice compared to the ones from young mice, we found a significant decrease in the number of hotspots 16 hours after transfection in aged neurons (**Figure 3A, C & D**). As a control, we used anisomycin to inhibit protein synthesis and confirmed a reduction in EGFP spots upon transfection with EGFP mRNA, confirming that the detected EGFP signal is due to the translation of the transcript (**Figure 3B, C & D**).

**Figure 3.**
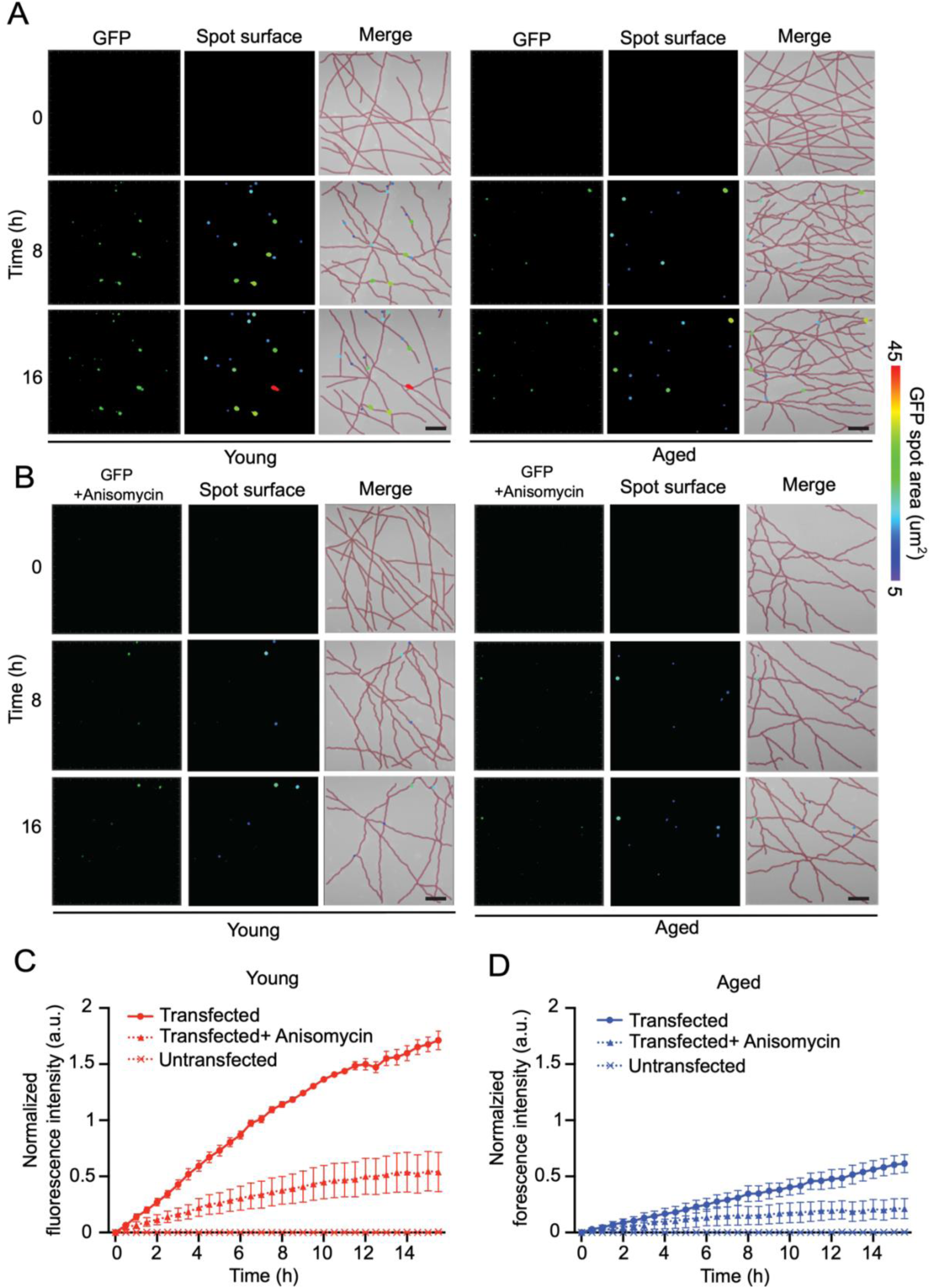
Impairement of axonal translation in aged neurons. (A) Representative images of translated GFP foci accumulating over 16 hours after transfection in DRG neuron axons from young and aged mice cultured in MFCs. (B) Representative images of translated GFP foci accumulation overtime in DRG neurons from young and aged mice treated with 40 μM anisomycin. For (A,B), left panels show foci of translated GFP (green), middle panels show GFP translational foci pseudocolored accorrding to their area (rainbow), right panels show merged images of indentified translated GFP foci masks with manually traced axons (red). Scale bar =15 µm, color bar represents the area of GFP spots. (C, D) Quantification of GFP translational foci fluorescence intensity over time in the experiments described in A and B.

### Proteomics analysis revealed differences in the axonal translatome of aged mice

Having observed differences in local axonal protein synthesis between aged sensory neurons and young ones, we set to explore the identity of the translated proteins. We made use of trans-well cultures to achieve physical separation of cell bodies and axons, and took advantage of O-propargyl-puromycin (OPP), an analog of puromycin, which can be incorporated in nascent polypeptides and CLICKED to biotin-azide (*21*, *33*). OPP-labelled proteins can then be retrieved with Streptavidin beads and identified through Mass Spectrometry (*21*, *33*). Proteomics analysis of the axonal and cell body translatomes revealed significant differences in the translational profiles of DRG neurons from aged and young mice both in axons (**Figure 4A & B**) and cell bodies (**Figure S5A & B**). GO categories analysis using the DAVID software suite showed an enrichment of proteins involved in axonal extension, cytoskeletal regulation and molecular transport in axons of young neurons (**Figure 4D**). Cell bodies of young neurons were enriched in proteins positively regulating translation (**Figure S5D**). Both aged axons and cell bodies displayed an enrichment in proteins involved in protein ubiquitination, retrograde transport, membrane fusion and response to hypoxia (**Figure 4C & Figure S5C**).

**Figure 4.**
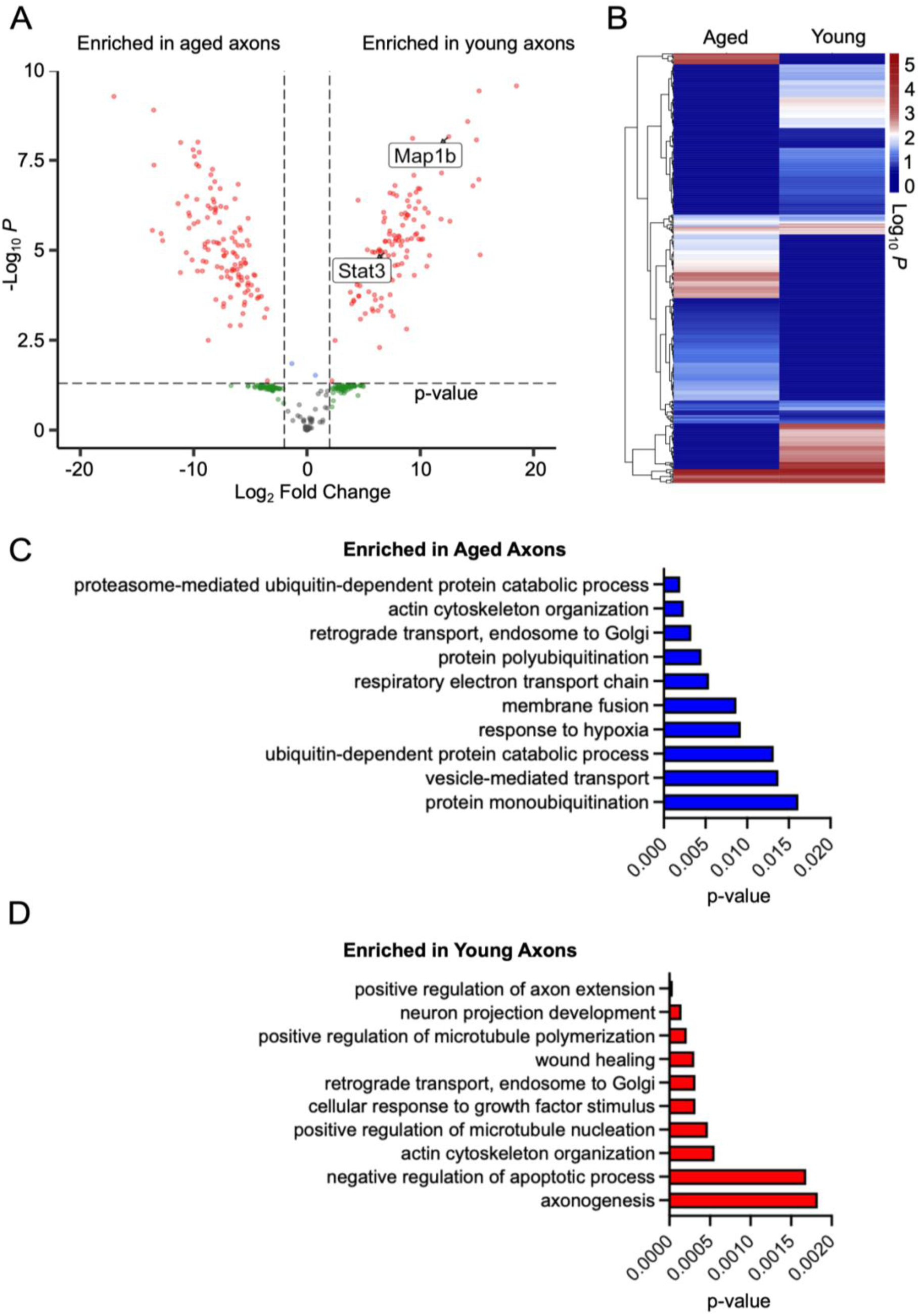
Mass spectrometry analysis of newly translated proteins in axons of DRG young and aged DRG neurons. (A) Volcano plot of the Mass Spectrometry (MS) analysis of axonal OPP-biotin labelled proteins enriched in young and aged mice. Vertical and horizonal dashed line mark the two-log fold change (log2(2) = 1) threshold and p value of 0.05, respectively. Red dots indicate candidates which pass both threshold, blue dots indicate candidates which does not pass the two-log fold threshold, green dots represent candidates which does not pass the p-value threshold. (B) Heat map of OPP-biotin labelled axonal proteins in DRG neurons from young and aged mice. (C, D) Gene ontology (GO) analysis of the molecular function of axonal proteins enriched in DRG neurons from aged and young mice, respectively.

After careful evaluation of all candidates, we decided to concentrate on the differences in the axonal translatome and to further characterize the Signal Transducer and Activator of Transcription 3 (STAT3) and the Microtubule-Associated Protein 1B (MAP1B), which were significantly enriched in axons of DRG neurons from young mice (**Figure 4A**). Indeed, both STAT3 and MAP1B are involved in axonal extension and microtubule stabilization (*34–37*), which have been reported to decrease during aging (*30*, *38*). Direct visualization of newly synthesized STAT3 and MAP1B by combining puromycin labeling and proximity ligation assay (PLA) as described by Schuman’s laboratory (*39*), revealed a significant decrease in translation of STAT3 and MAP1B in axons of aged neurons compared to young ones (**Figure 5A & B, Figure 5C & D**), but not in cell bodies (**Figure 5A & C, Figure S6C, and Figure S6D**). We verified that the observed PLA signal came from protein translation by using anisomycin to inhibit protein synthesis (**Figure S6A-D & Figure 5B, D**).

**Figure 5.**
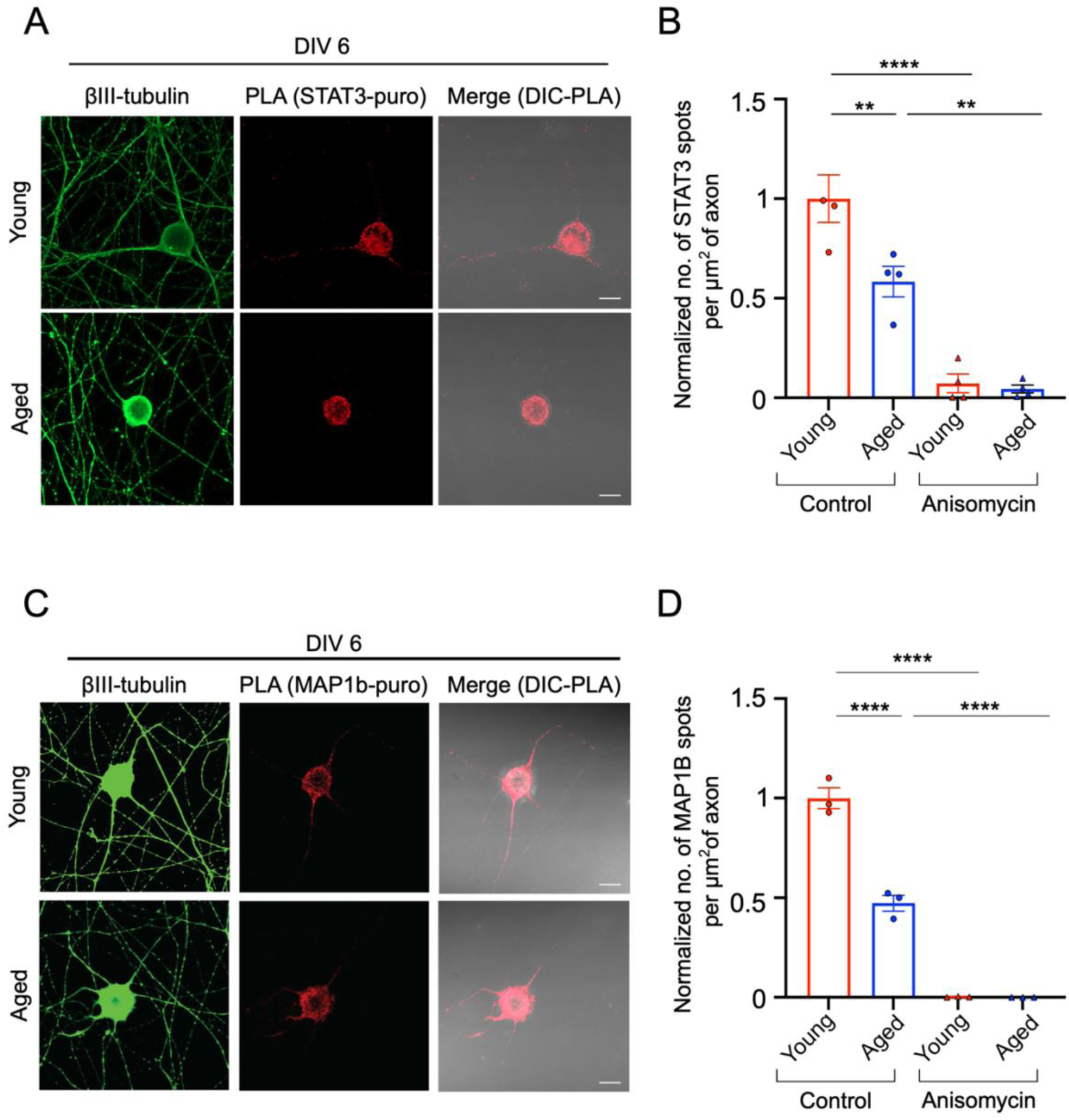
STAT3 and MAP1B axonal protein translation in aged mice at DIV 6. (A) Representative images of newly synthezied STAT3 protein in puromycin-treated DRG neurons labelled with βIII-tubulin (green) detected by PLA between α-puromycin and α-Stat3 antibodies (red). Neurons were incubated with puromycin for 15 minutes to label newly synthesized protein. Scale bar = 10 µm. (B) Quantification of axonal puro-PLA signal from experiments repesented in (A) (n= 40 cells from 4 biological replicates; means ± SEM; ns, not signifcant; \**p* < 0.05 ; one-way *ANOVA*). (C) Representative images of newly synthezied MAP1B protein in puromycin-treated DRG neurons labelled with βIII-tubulin (green) detected by PLA between α-puromycin and α-Map1b antibodies (red). Neurons were incubated with puromycin for 15 minutes to label newly synthesized protein. Scale bar = 10 µm. (D) Quantification of axonal puro-PLA signal from experiments described in (C) (n= 30 cells from 3 biological replicates; means ± SEM;\*\*\*\**P* < 0.0001 ; one-way *ANOVA*).

### STAT3 and MAP1B mRNA accumulate in RNA granules of aged axons

We performed proteomics analysis from neurons cultured in filter chambers at DIV 6 to minimize the glial contamination on the axonal size. Our previous experiments in MFCs were performed at DIV 14 to allow sufficient time for the axons to cross to the axonal site of the chamber. To support the results of the proteomics analysis, we assessed cytoplasmic viscosity in axons (**Figure S7A, B & C**) and cell bodies (**Figure S7D, E & F**) of aged neurons at DIV 6 in culture and found it to be reduced compared to young ones consistent with our previous findings at DIV 14 (**Figure 1F, G & H, and Figure S2 A, B & C**).

G3BP1 and FMRP proteins, which associate with RNA granules (*8*, *9*, *27*), have been shown to undergo LLPS. Interestingly, dissolution of RNA granules might enhance the rate of regeneration by promoting axonal translation (*8*, *9*, *27*). We tested whether the decrease of ATP concentration and increase in cytoplasmic viscosity observed in aged axons at DIV 6, also influences the dissolvement of RNA granules, and subsequently the release of mRNAs. We previously showed that MAP1B mRNA is associated with axonal FMRP granules in DRG neurons (*8*). In this study, using RNAscope coupled with immunostaining, we were able to detect STAT3 association with axonal G3BP1 granules (**Figure 6A**). We observed a significant increase in the overall axonal STAT3 mRNA levels and the number of axonal G3BP1 granules positive for STAT3 mRNA in aged neurons compared to young ones (**Figure 6A & B**). Similarly, we identified a significant increase in axonal MAP1B mRNA levels and number of FMRP granules positive for MAP1B mRNA in aged neurons compared to young ones (**Figure 6C & D**). Taken together, these findings suggest there was an increase in the cytoplasmic viscosity of axons from aged mice, which subsequently impaired the solubilization of both G3BP1 and FMRP RNA granules and prevented the release of STAT3 and MAP1B mRNA.

**Figure 6.**
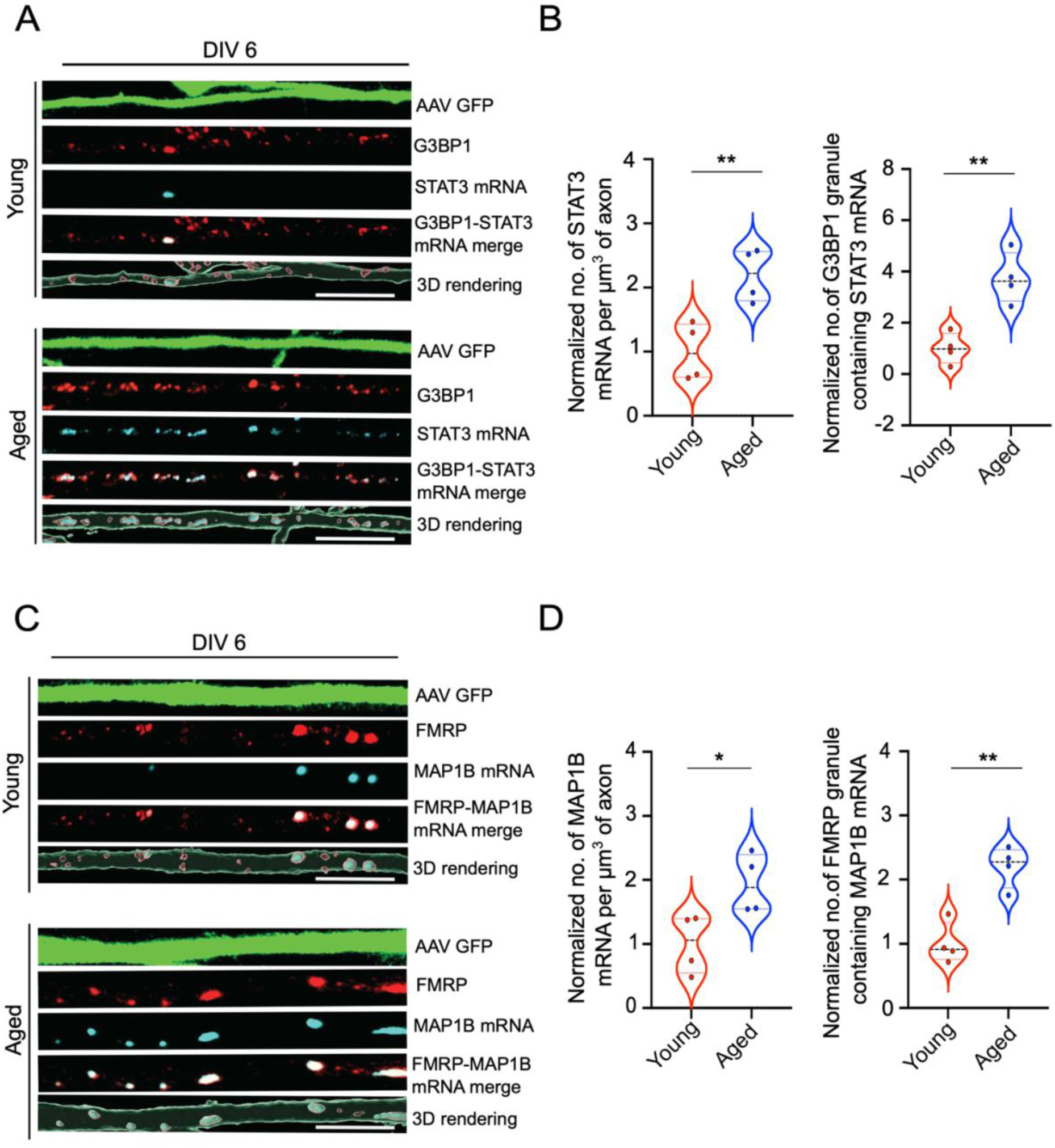
STAT3 and MAP1B association to G3BP1 and FMRP granules in axons of young and aged DRG neurons at DIV 6. (A) Representative image of integrated in situ hybridization (ISH) for STAT3 mRNA (red) and G3BP1 RNA granule (turquoise) immunostaining in DRG neurons from young and aged mice. Neurons are labeled with GFP AAV (green). Last panels show a 3D representation of STAT3 mRNA inside G3BP1 granules along the axon. Scale bar = 5 µm. (B) Quantification of axonal STAT3 mRNA inside axonal G3BP1 granules and number of G3BP1 granules containing STAT3 mRNA as described in experiment (A) (n= 32 cells from 4 biological repeats; means ± SEM; \*\**p* < 0.01 ; unpaired *t* test). (C) Representative image of integrated in situ hybridization (ISH) for MAP1B mRNA (red) and FMRP RNA granule (turquoise) immunostaining in DRG neurons from young and aged mice. Neurons are labeled with GFP AAV (green). Last panels show a 3D representation of MAP1B mRNA inside FMRP granules along the axon. Scale bar = 5 µm. (D) Quantification of axonal MAP1B mRNA inside axonal FMRP granules and number of FMRP granules containing MAP1B mRNA as described in experiment (A) (n= 32 cells from 4 biological replicates; means ± SEM; \**p* < 0.05, \*\**p* < 0.01 ; unpaired *t* test).

### NMN treatment increases cytosolic fluidity and decreases G3BP1 and FMRP RNA granule aggregation in aged neurons

As previously discussed, our findings showed an accumulation of STAT3 and MAP1B mRNA inside G3BP1 and FMRP RNA granules, respectively, in axons from aged mice at DIV 14 (**Figure S8A & B, Figure S8C & D**). We also observed a decline in axonal translation using both puromycin labelling (**Figure S3C & D**) and live cell imaging of transfected EGFP mRNA translation (**Figure 3**). We recently showed that a chronic 7 days treatment with Nicotinamide Mononucleotide (NMN), a precursor of NAD^+^, can increase mitochondrial activity and ATP levels in ALS motor neurons (MN), leading to a fluidification of their axonal cytoplasm (*11*). In addition, NMN supplementation has been shown to play a key role in improving mitochondrial health during aging and is linked to improved survival of neurons in degenerative disease (*40–42*). Thus, we wondered if NMN treatment would be able to increase intracellular ATP levels in aged sensory axons and, in turn, enhance axonal translation.

Similarly to we found in MNs (*11*), a week of incubation with NMN was able to increase intracellular ATP levels in DRG neurons from aged mice (**Figure S9A**). Further, we observed a decrease in axonal cytoplasmic viscosity of aged mice after NMN treatment (**Figure S9B & C**). NMN treatment of ALS MNs was able to dissolve TDP-43 axonal aggregates (*11*). Likewise, the volume of G3BP1 positive granules (**Figure 7A & B, and Figure S9D**), but not their number (**Figure 7A & B**), was significantly decreased in axons of aged DRG neurons after NMN treatment. Interestingly, the number of FMRP positive granules (**Figure 7C & D**), but not their volume (**Figure 7C & D, and Figure S9E**), was significantly decreased in axons of aged DRG neurons after NMN treatment. In accordance to what we observed in the disease model, where MNs derived from healthy individual did not show changes after NMN treatment (*11*), we did not see any effect of NMN treatment on ATP levels (**Figure S9A**), cytoplasmic fluidity (**Figure S9B & C**) and G3BP1/FMRP positive granules in young neurons (**Figure 7A – D, Figure S9D & E)**.

**Figure 7.**
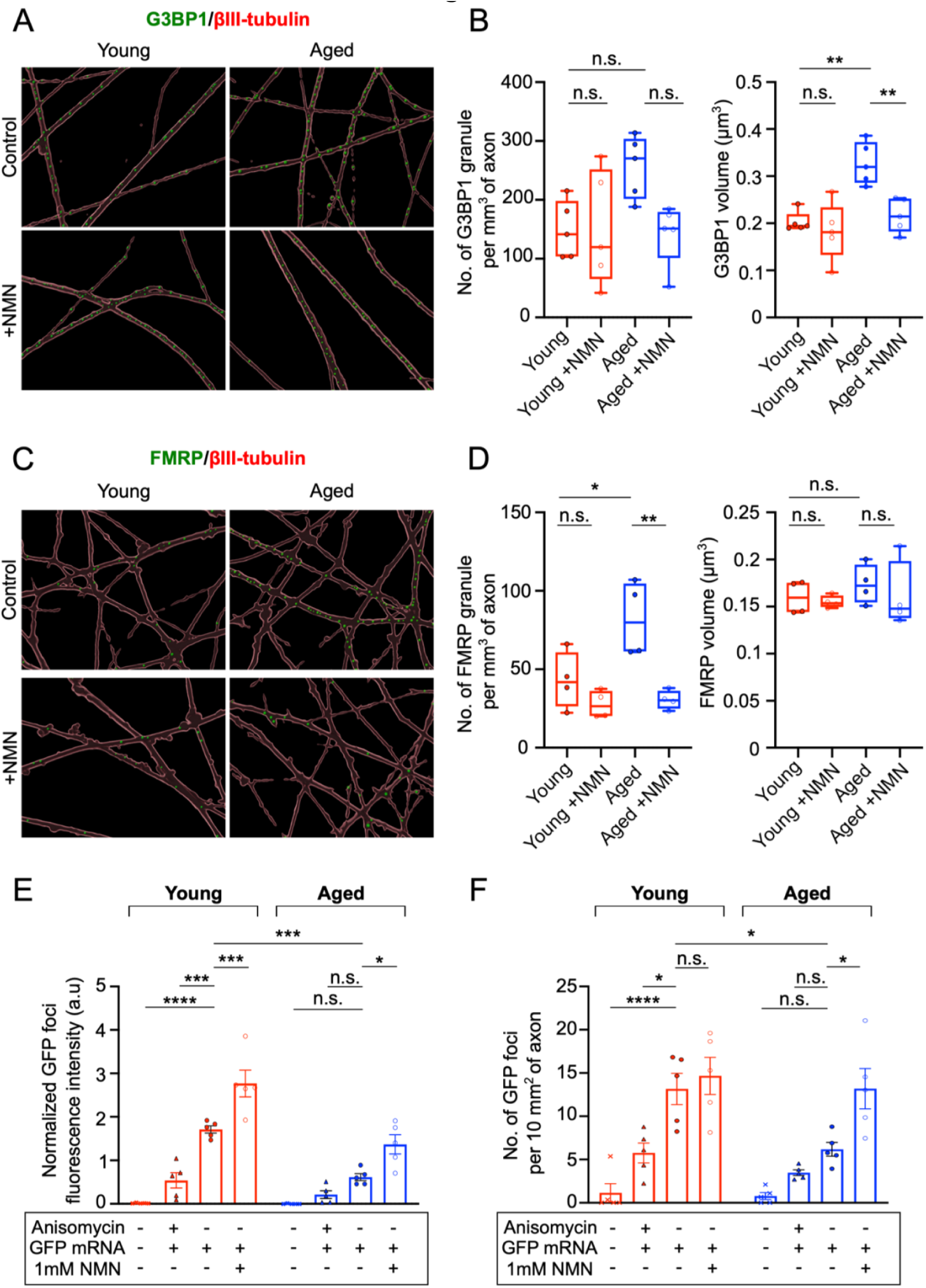
NMN treatment increases axonal translation in aged neurons. (A) 3D surface rendering of axonal G3BP1 granules (green) labelled with βIII-tubulin (red) of young and aged mice, with and without NMN treatment. (B) Quantification of number and volume of axonal G3BP1 RNA granules of young and aged neurons, with and without NMN treatment, as described in (A) (n= 5; means ± SEM; ns, not significant; \**p* < 0.05, \*\**p* < 0.01; one-way *ANOVA*). (C) 3D surface rendering of FMRP granules (green) along the axons labelled with βIII-tubulin (red) of young and aged mice, with and without NMN treatment. (D) Quantification of number and volume of axonal FMRP RNA granule of young and aged neurons, with and without NMN treatment, as described in (C) (n= 4; means ± SEM; ns, not significant; \**p* < 0.05, \*\**p* < 0.01; one-way *ANOVA*). (E) Quantifcation of GFP translational foci fluorescence intensity in DRG neurons from young and aged mice, with and without NMN treatment, following a 16-hour incubation period (n= 5; means ± SEM; ns, not significant; \**p* < 0.05, \*\*\**p <* 0.001,\*\*\*\**p* < 0.0001; one-way *ANOVA*). (F) Quantifcation of the number of GFP translational foci in DRG neurons from young and aged mice, with and without NMN treatment, following a 16-hour incubation period (n = 5; means ± SEM; ns, not significant; \**p* < 0.05, \*\*\*\**p* < 0.0001; one-way *ANOVA*).

### NMN treatment increases axonal translation in aged neurons

Dissolution of G3BP1 RNA granules has been associated with increased translation (*9*). In a previous work we also showed that FMRP axonal granules can restrict the translation of associated mRNAs, such as MAP1B (*8*). Thus, we tested axonal translation levels in young and aged sensory neurons after NMN treatment. We transfected the axons of young and aged DRG neurons in MFCs, as previously described and recorded the increase of axonal GFP protein for 16 hours after transfection. Using this method, we observed an increase in axonal translation in aged neurons, as indicated by elevated level of fluorescence intensity and the number of translation-associated EGFP foci following NMN supplementation (**Figure 7E & F, Figure S10A**). Interestingly, we observed an increase in the fluorescence intensity of EGFP translation foci following NMN treatment in both young and aged axons (**Figure 7E & D**), while the number of axonal translational foci increased only in aged axons (**Figure 7F**). NMN treatment was also unable to increase the area of axonal EGFP translational foci in both young and aged neurons (**Figure S10A-C**).

In addition to the effect of NMN treatment on the overall levels of axonal translational activity, we observed increased axonal translation of STAT3 and MAP1B in aged neurons following NMN supplementation (**Figure 8A-D**), which was not observed in the cell body (**Figure 8A & B, Figure S11C & D**). Anisomycin was used as a negative control to confirm that the Puro-PLA signal observed relates to translational activity (**Figure S11A-D & Figure 8C-D**).

**Figure 8.**
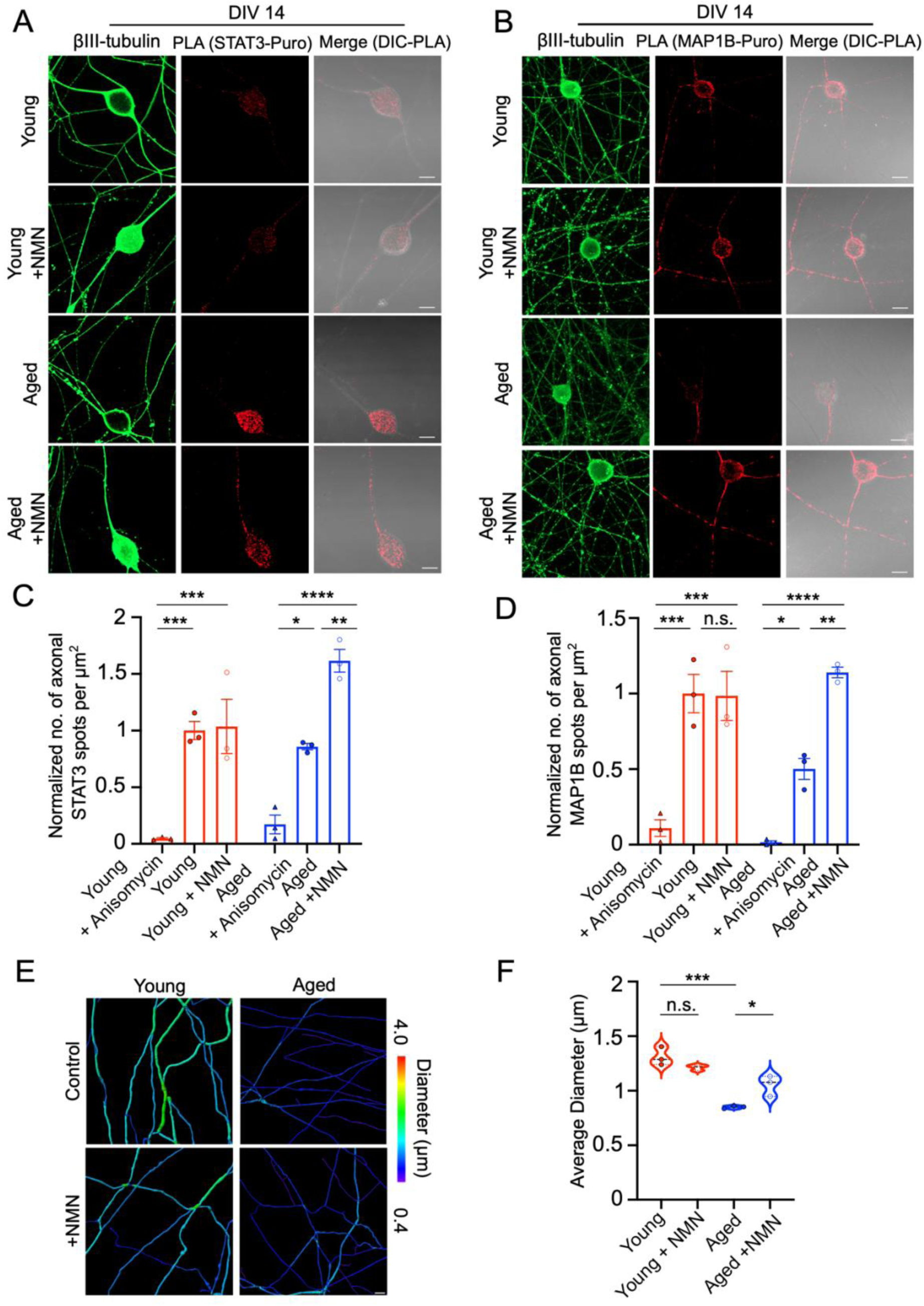
NMN treatment increases axonal translation of STAT3 and MAP1B in aged neurons. (A) Representative images of newly synthezied STAT3 protein in puromycin-treated DRG neurons cultured at DIV 14, labelled with βIII-tubulin (green) and detected by PLA between α-puromycin and α-Stat3 antibodies (red), with and without NMN treatment. Neurons were incubated with puromycin for 15 minutes to label newly synthesized protein. Scale bar = 10 µm. (B) Representative images of newly synthezied MAP1B protein in puromycin-treated DRG neurons cultured at DIV 14, labelled with βIII-tubulin (green) and detected by PLA between anti-puromycin and anti-Map1b antibodies (red), with and without NMN treatment. Neurons were incubated with puromycin for 15 minutes to label newly synthesized protein. Scale bar = 10 µm. (C) Quantification of translated axonal STAT 3 as described in (A) (n= 30 cells from 3 biological repeats; means ± SEM; \**p* < 0.05, \*\**p* < 0.01, \*\*\**p* < 0.001, \*\*\*\**p* < 0.0001; one-way *ANOVA*). (D) Quantification of translated axonal Map1B as described in (B) (n= 30 cells from 3 biological repeats; means ± SEM; ns; not significat; \**p* < 0.05, \*\**p* < 0.01, \*\*\**p* < 0.001, \*\*\*\**p* < 0.0001; one-way *ANOVA*). (E) 3D surface rendering of the axonal diameter of young and aged mice, with and without NMN treatment. Scale bar = 5 µm, color bar represents the axonal diameter. (F) Quantification of average axonal diamater as described in (E) (n= 3; means ± SEM; \**p* < 0.05, \*\*\**p* < 0.001; one-way *ANOVA*).

MAP1B translation in DIV 14 neuronal culture mirrored the findings at DIV 6, showing a significant reduction in axonal translation in aged neurons compared to young neurons (**Figure 8B & D, Figure S11F**). In contrast, STAT3 axonal translation did not exhibit a significant decrease in aged neurons (**Figure 8A & C**). However, when normalizing axonal STAT3 translation to total translational activity of both axons and cell bodies, axonal translation remained lower in aged neurons compared to young neurons (**Figure S11E**). MAP1B plays a critical role in cytoskeleton assembly, a process essential for neuronal growth (*43*). Similarly, STAT3 has been shown to interact with other proteins such as stathmin to promote microtubule stabilization (*37*). In *Drosophila*, MAP1B helps has been implicated in the regulation of axon diameter (*44*) and MAP1B-deficient mice exhibit a smaller axonal diameter, which has been linked to impaired sciatic nerve motor function (*45*). Furthermore, STAT3 might influence axonal caliber by mediating IL-6 signaling within the axon, as IL-6 deficiency has been reported to cause microtubule destabilization and smaller axonal caliber (*46*). Interestingly, in this study we found that axons of aged neurons have a smaller diameter compared to ones of young neurons (**FIGURE 8E & F**). In accordance with our data on translation, NMN treatment was able to increase axonal diameter in aged neurons (**FIGURE 8E & F**).

In summary, we demonstrated that NMN treatment was able to rescue ATP levels, cytosol viscosity, axonal translation and axon diameter in aged neurons.

## Discussion

Aging is characterized by the loss of several cellular and molecular functions, increasing the body’s vulnerability to metabolic and neurodegenerative diseases (*47–49*). Decline in the regenerative capacity of neurons has also been described as one of the consequences of aging (*30*, *50*), and axonal translation is a critical factor which influence the efficiency of neuronal regeneration (*9*, *30*). Axonal protein synthesis is an energy-dependent process and thus sensitive to changes in ATP availability in the cellular environment. Mitochondria play a key role in ATP supply to axons (*51*). A decline in mitochondrial trafficking has been reported in peripheral neurons of 2 year old aged mice (*17*), as well as other aging organisms (*13*, *15*). We showed that a loss of motile mitochondria can already be observed in peripheral neurons of aged mice (1 year old). We also found that the number of active mitochondria was reduced, leading to a subsequent decline in intracellular ATP levels (**Figure 1 A-D, Figure S1A-E**).

Our FRAP analysis highlighted how the reduced ATP levels observed in aged neurons are associated with a decrease in the cytosolic fluidity of both sensory neuron axons and cell bodies, though more prominent in axons (**Figure 1F, G & H, Figure S2A, B & C**). This finding aligns with emerging evidence that ATP levels can modulate protein aggregation, both *in vitro* and *in vivo* (*11*, *26*, *52*, *53*). Recently, we reported that in human motor neurons derived from human pluripotent stem cells (hiPSC), cytosolic ATP levels affect axoplasmic viscosity and the formation of TDP-43 protein aggregates in Amyotrophic Lateral Sclerosis (ALS) (*11*). In the current study, we showed that the decrease cytosolic fluidity of aged sensory axons modulates G3BP1 and FMRP RNA granules, preventing their dissolution and the release of their sequestered mRNA. In addition, G3BP1-positive RNA granules have been shown to be dissolved following neuronal injury, a process believed to facilitate the release and translation of sequestered mRNAs (*9*). In addition, the results of our FRAP analysis align with the recent findings reported using rheology analysis of genetically encoded multimeric nanoparticles (GEMs) in 2 year old sensory neurons, which showed an increased viscosity in both axons and cell bodies (*17*), supporting our hypothesis of an impairment of ATP-dependent viscoadaption of the cytosol of aged DRG neurons. However, using a different method to evaluate viscosity, time-resolved anisotropy analysis, the same study also reported a specific increase in the cytosolic viscosity of the cell body, but not axons, of aged neurons (*17*). These discrepancies might be the result of the limitations of the different techniques used to evaluate cytosolic viscosity, as well as the regions of measurement in axons, the maturation of the axonal network, or the overall aged of the animals used in the studies, with their mice being two years old, while our aged mice are 52-63 weeks old.

Using microfluidic chambers, we were able to document a decline in axonal translation in aged axons. Proteomics analysis further revealed age-dependent changes in protein expression, in agreement with what was reported for mature CNS neuronal culture (*30*). Axons of sensory neurons isolated from young mice synthesized more proteins involved in cytoskeleton organization, axonal outgrowth and protein synthesis. In contrast, axons of aged neurons showed an enrichment of proteins associated with stress-response and protein turnover. Notably, STAT3 and MAP1B were among the proteins whose levels were negatively altered in aged axons. STAT3 is a key player in axonal regeneration (*2*), while MAP1B is important during development for axonal growth and guidance (*19*). In addition to its role in development, MAP1B levels remain elevated in adult DRG neurons, where it is responsible for axonal turning and branching (*19*). Based on our data, we speculate that the increase in cytosol viscosity caused by reduced ATP levels negatively impacts the release of STAT3 and MAP1B mRNAs from G3BP1 and FRMP granules and subsequently their availability for translation. Interestingly, a recent paper using *Xenopus* egg extracts described how higher cytoplasmic viscosity due to increased protein concentration has a negative effect on translational activity (*32*).

Importantly, supplementation with NMN, a percussor of NAD^+^, effectively increased intracellular ATP levels in cultured DRG neuron axons and promoted the dissolution of both G3BP1 and FMRP granules in aged neurons, thereby facilitating the release of STAT3 and MAP1B mRNA and restoring their axonal translation levels. Indeed, NMN has been reported to mitigate age-related decline, increase energy production and neuronal survival in mice (*40*, *54*, *55*), and reduce pathological aggregation of proteins in ALS (*11*). Interestingly, the effects of NMN on STAT3 and MAP1B translation was specific to the axon, as no significant increase was observed in the cell body. In our proteomics screen we also found upregulated in young axons translation initiation factors, such as EiF2, and ribosomal proteins, which could constitute a parallel mechanism to the ATP-mediated regulation of granule formation, which mediates the higher axonal translation rate of young neurons compared to aged ones.

These results suggest that targeting energy metabolism and RNA granule dynamics could potentially enhance axonal translation and subsequently improve axonal health during aging in the nervous system.

Taken together, our findings highlight how mitochondrial dysfunction in aged neurons and consequent lower ATP levels, negatively impacts axonal translation. We believe that axonal ATP plays an important regulatory role in the assembly and dissolution of RNA granules, thereby regulating the accessibility for translation of their associated mRNAs. Thus, our work expands the understating on how ATP affects axonal translation during aging in sensory neurons, offering new insight into potential therapeutic strategies for improving axonal function and regeneration.

## Acknowledgments

We thank Prof. Mike Fainzilber and Prof. Giampietro Schiavo for critical reading of the manuscript. We also thank OIST animal facility for taking care of our experimemtal animals and the procurement of aged mice. We would also like to thank OIST Instrumental Analysis section and in particular Dr. Yoshitoshi Hirao for helping us with the MS run.

## Contributions

M.F.E., L.G. and M.T. conceived the project, performed the experiments, collected the data, and drafted the manuscript. Y.A. and S.B. optimized the OPP clicking protocol. S.D.F.R. helped with the mass spectrometry data. R.A. and M.E.R. designed the algorithm for the identification of GFP translational spots. S.D.F.R., R.A., Y.A., S.B., T.H.T., M.E.R. helped in the development of ideas. T.H.T. performed preliminary proof of concept axperiments. M.F.E., L.G. M.C. and M.T. participated in the editing of manuscript.

## Funding

M.T. and M.E.R. acknowledge the internal funding from the Okinawa Institute of Science and Technology Graduate University and JSPS/Kakenhi C Research Grant (#23K27107) in the current study. L.G. was supported by JSPS/Kakenhi C research Grants (#21K06400).

## Declaration of conflict of interest

The authors declared no potential conflicts of interest with respect to the research, authorship, and/or publication of this article.

## Notes

### Competing Interest Statement

The authors have declared no competing interest.

