## Supplementary Figures and Figure Legends for "Impairment in axonal translation and cytoplasmic viscosity during aging in sensory neurons"

**Title:**

#### Supplementary Figure Legends

**Figure S1. Mitochondrial motility in DRG neurons from young and aged mice.** (A) Representative images of axonal mitochondria labelled with MitoTracker® (green) from young and aged mice (n= axons from 5 biological replicates; means ± SEM; ns, not signifcant; unpaired *t* test). Scale bar = 10μm. (B) Kymograph representation of axonal mitochondrial motility in DRG neurons from young and aged mice. (C, D) Quantification of the frequency of motile axonal mitochondria observed inside microfluidic chambers in both anterograde and retrograde direction, respectively. (E) Quantification of the total displacement of motile axonal mitochondria in anterograde (left) and retrograde (middle) direction (n= 6; dashed line on mean; ns, not signifcant; *t* test), as well as the percentage of motile axonal mitochondria (right) between DRG neurons from young and aged mice (n young= 29 carriers, n aged= 26 carriers; dashed line on mean; ****$p < 0.0001$; unpaired *t* test).

**Figure S2. Cell body of aged neurons at DIV14 display decreased citoplamisc viscosity compared to young ones.** (A) Cell body of DRG neurons neurons expressing cGFP (pseudo-color) at DIV 14 from young and aged mice. Panels from left to right show the FRAP ROI before photo-bleaching, during bleaching, and post recovery. Scale bar = 5 μm. The color bar represents cGFP fluorescence intensity. (B) Fluorescence intensity recovery profile of DRG neurons cell bodies at DIV 14 from young and aged mice. Mean ± 95% CI. (C) Mobile cGFP fraction representing cytosolic fluidity estimation from the last 20 seconds of the experiment described in (A) (young n= 28, aged n= 24 cell bodies from 3 biological repeats; means ± SEM; **$p < 0.01$; unpaired t test).

**Figure S3. Puromycinilation of newly synthesized protein in MFCs.** (A) Representative images of a time-course of puromycinilation of newly synthesized axonal proteins in MFCs. DRG neurons were incubated with puromycin for 15 minutes before being washed and fixed at 0, 30 and 60 minutes after the removal of puromycin. Scale bar = 100μm. (B) Quantification of puromycin fluorescence intensity in axons overtime as described in (A) (n = axons from 4 biological replicates; dashed line at mean; *$p < 0.05$,

**p < 0.01 ; one-way *ANOVA*). (C) Representative images of newly synthesized protein labelled using puromycin (green) in DRG neurons axons of young and aged mice labelled with βIII-tubulin (orange). Puromycin signal was pseudo-colored accordingly to the fluorescence intensity (rainbow, right panel). Scale bar = 20 μm. (D) Quantification of puromycin fluorescence intensity of the experiments described in (C) (n= axons from 4 biological replicates; dashed line on mean; ** $P < 0.01$, unpaired *t* test).

**Figure S4. Outline of GFP translational foci intensity quantification.** (A) Represenative images representing the workflow of the Phyton script for the quantification of the intensity of axonal GFP translational foci. Scale bar = 20μm. (B) Schematic workflow of the algorithm underlying Python script desibed in (A).

**Figure S5. Mass Spectrometry analysis of newly translated proteins in the cell body of DRG neurons from young and aged mice.** (A) Volcano plot of the Mass Spectrometry (MS) analysis of OPP-biotin labelled proteins enriched in the cell body of young and aged neurons. Vertical and horizonal dashed line marked threshold of two log fold change ($\log_2(2) = 1$) and p value of 0.05, respectively. Red dots indicate candidates passing both thresholds, blue dots indicate candidates which does not pass the two-log fold threshold, green dots represent candidates which does not pass the p-value threshold. (B) Heat map of OPP-biotin labelled proteins in the cell body of DRG neurons from young and aged mice. (C, D) Gene ontology (GO) analysis of the molecular function of proteins that are enriched in the cell body of DRG neurons from aged and young mice respectively.

**Figure S6. STAT3 and MAP1B protein translation in the cell body of young and aged DRG neurons at DIV 6.** (A) Representative images of newly synthesized STAT3 protein in anisomycin-treated DRG neurons labelled with βIII-tubulin (green) detected by PLA between anti-puromycin and α-Stat3 antibodies (red). Neurons were incubated with anisomycin for 30 minutes prior to puromycin incubation for 15 minutes. Scale bar = 10 μm. (B) Representative images of newly synthesized MAP1B protein in anisomycin-treated DRG neurons labelled with βIII-tubulin (green) detected by PLA between α-puromycin and α-Map1B antibodies (red). Neurons were incubated with anisomycin for

30 minutes prior to puromycin incubation for 15 minutes. Scale bar = 10 μm. (C) Quantification of puro-PLA signal in the cell body from experiments described in **Figure 5A** (n= 40 cells from 4 biological repeats; means ± SEM; ns, not signifcant; *$p$ < 0.05, **$p$ < 0.01; one way *ANOVA*). (D) Quantification of puro-PLA signal in the cell body from experiments described in **Figure 5C** (n= 40 cells from 4 biological replicates; means ± SEM; ns, not signifcant; ****$p$ < 0.0001; one way *ANOVA*).

**Figure S7. Cell body of aged neurons at DIV6 display increased cytoplasmic viscosity compared to young ones.** (A) Axon of DRG neurons expressing cGFP (pseudo-color) from young and aged mice. Panels from left to right show the FRAP ROI before photo-bleaching, during bleaching, and post recovery. Scale bar = 5 μm. The color bar represents cGFP fluorescence intensity. (B) Axonal fluorescence intensity recovery profile of DRG neurons from young and aged mice. Mean ± 95% CI. (C) Mobile cGFP fraction representing cytosolic fluidity estimation from the last 20 seconds of the experiment described in (A) (n= 21 axons from 3 biological repeats; means ± SEM; ****p < 0.0001; unpaired t test). (D) Cell body of DRG neurons from young and aged mice expressing cGFP (pseudo-color). Panels from left to right show ROI before photo-bleaching, during bleaching, and post recovery. Scale bar = 5 μm. The color bar represents cGFP fluorescence intensity. (E) Fluorescence intensity recovery profile of DRG neurons cell body from young and aged mice. Mean ± 95% CI. (F) Mobile cGFP fraction representing cytosolic fluidity estimation from the last 20 seconds of the experiment described in (D) (young n= 29, aged n=30 cell bodies from 3 biological repeats; means ± SEM; **p < 0.01 ; unpaired t test).

**Figure S8. Axonal STAT3 and MAP1B mRNAs in young vs aged neurons at DIV 14.** (A) Representative images of integrated *in situ* hybridization (ISH) for STAT3 mRNA (red) and G3BP1(turquoise) immunostaining in the axon of young and aged mice. Neurons are labeled with GFP AAV (green). Last panels show a 3D representation of STAT3 mRNA inside axonal G3BP1 granules. Scale bar = 5 μm. (B) Quantification of axonal STAT3 mRNA inside axonal G3BP1 granules and number of G3BP1 granules containing STAT3 mRNA as described in (A) (n= 32 neurons from 4 biological replicates; means ± SEM; *$p$

< 0.05,***p* < 0.01; unpaired *t* test). (C) Representative images of integrated *in situ* hybridization (ISH) for MAP1B mRNA (red) and FMRP (turquoise) immunostaining in DRG neurons from young and aged mice. Neurons are labeled with GFP AAV (green). Last panels show a 3D representation of MAP1B mRNA inside axonal FMRP granules. Scale bar = 5 μm. (D) Quantification of axonal MAP1B mRNA inside axonal FMRP granules and number of FMRP granules containing MAP1B mRNA as described in (C) (n= 32 cells from 4 biological experiments ; means ± SEM; **p* < 0.05,***p* < 0.01; unpaired *t* test). (E) Representative images of integrated *in situ* hybridization (ISH) with negative and positive control RNA Scope probes (turquoise). Neurons are labeled with GFP AAV (green). Scale bar = 5 μm.

**Figure S9. Effects of NMN treatment on intracellular ATP level and RNA granule condensation in young and aged neurons.** (A) *In vitro* bioluminescence measurement of intracelllular ATP levels in DRG neurons from young and aged mice with and without NMN treatment (n = 3; means ± SEM; **p* < 0.05, ****p* < 0.001; one-way *ANOVA*). (B) Fluorescence intensity recovery profile of DRG neurons from young and aged mice with and without NMN treatment. Mean ± 95% CI. (C) Mobile cGFP fraction representing cytosolic fluidity estimation from the last 20 seconds of the experiment described in (B) (young n = 34, aged n = 33, young NMN n = 25, aged NMN n = 25 cells from at least 3 biological repeats; means ± SEM; ns, not significant; *****p* < 0.0001; one-way *ANOVA*). (D) Relative frequency of the volume of axonal G3BP1 granules in young and aged neurons, with and without NMN treatment, as described in Figure 7A (***p* < 0.01, ****p* < 0.001; two way ANOVA, Sidak comparison post-test). (E) Relative frequency of the volume of axonal FMRP granules in young and aged neurons, with and without NMN treatment, as represented in Figure 7C.

**Figure S10. Effect of NMN treatment on axonal translation in young and aged neurons.** (A) Representative images of translational GFP foci accumulation 16 hours after transfection in young and aged neurons. From top to bottom; neurons not transfected with GFP mRNA, neurons transfected with GFP mRNA and treated with anisomycin, neurons transfected with GFP mRNA, neurons transfected with GFP mRNA and treated with NMN. From left to right; translated GFP foci (green), translated GFP foci pseudo-colored according to their area (rainbow), merged images of translated GFP foci with manually traced axons (red). Scale bar = 15 µm, color bar represents translated GFP area. (B) Quantification of the distribution of translated GFP foci' area 16 hours after transfection in young and aged neurons, with and without NMN treatment as described in (A). (C) Quantification of average translated GFP foci' area following 16 hours transfection in young and aged neurons, with and without NMN treatment as described in (A) (young n=344, aged n=244, young NMN n=590, aged NMN n=618, axons from 5 biological repeats; means ± SEM; ns, not significant; ****$P < 0.0001$; two-way *ANOVA*).

**Figure S11. STAT3 and MAP1B protein translation in the cell body of young and aged neurons at DIV 14.** (A) Representative images of newly synthezied STAT3 protein in anisomycin-treated DRG neurons labelled with βIII-tubulin (green) detected by PLA between α-puromycin and α-Stat3 antibodies (red). Neurons were incubated with anisomycin for 30 minutes prior to puromycin incubation for 15 minutes. Scale bar = 10 µm. (B) Representative images of newly synthesized MAP1B protein in anisomycin-treated DRG neurons labelled with βIII-tubulin (green) detected by PLA between α-puromycin and α-Map1B antibodies (red). Neurons were incubated with anisomycin for 30 minutes prior to puromycin incubation for 15 minutes. Scale bar = 10 µm. (C) Quantification of puro-PLA signal in the cell body from experiments described in Figure 8A (n= 30 cells from 3 biological replicates; means ± SEM; ns, not signifcant; *$p < 0.05$; one-way *ANOVA*). (D) Quantification of puro-PLA signal in the cell body from experiments described in Figure 8B (n= 30 cells from 3 biological repeats; means ± SEM; ns, not signifcant; *$p < 0.05$, **$p < 0.01$; one-way *ANOVA*).(E) Quantification of axonal STAT3 translation ratio in DRG neurons of young and aged neurons (n= 30 cells from 3 biological repeats; means ± SEM; **$p < 0.01$; unpaired *t* test). (F) Quantification of axonal MAP1B translation ratio in DRG neurons of young and aged neurons (n= 30 cells from 3 biological repeats; means ± SEM; **$p < 0.01$; unpaired *t* test).

### Supplementary Figure 1

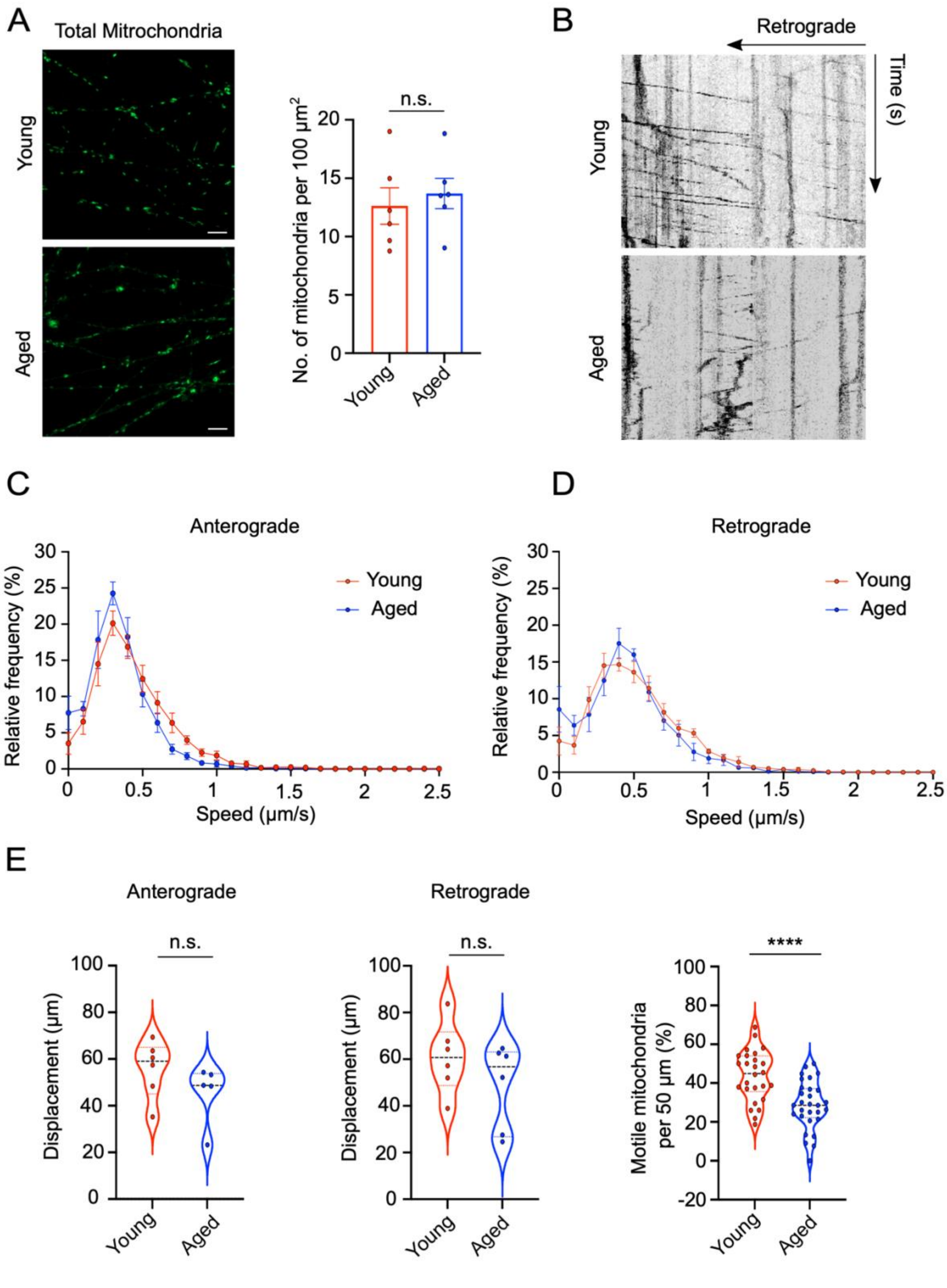

### Supplementary Figure 2

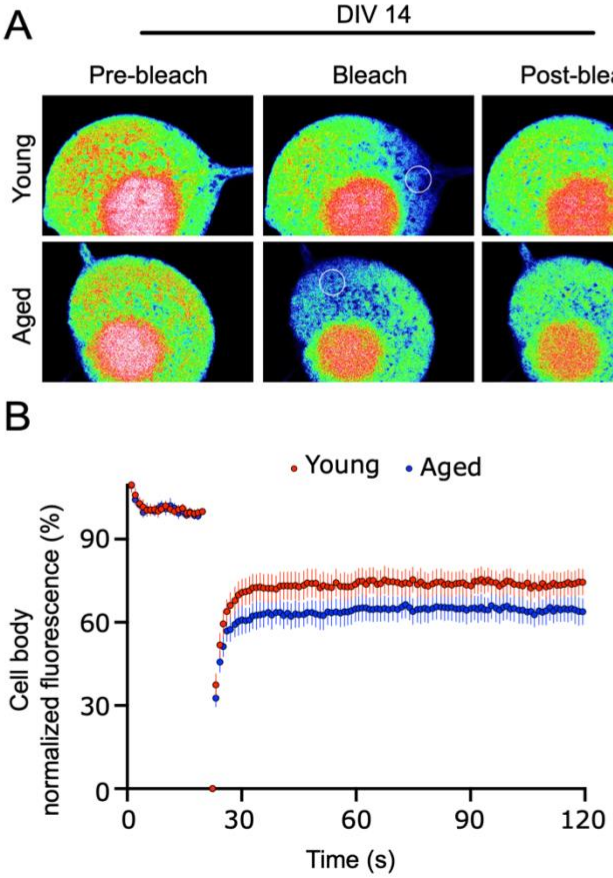

### Supplementary Figure 3

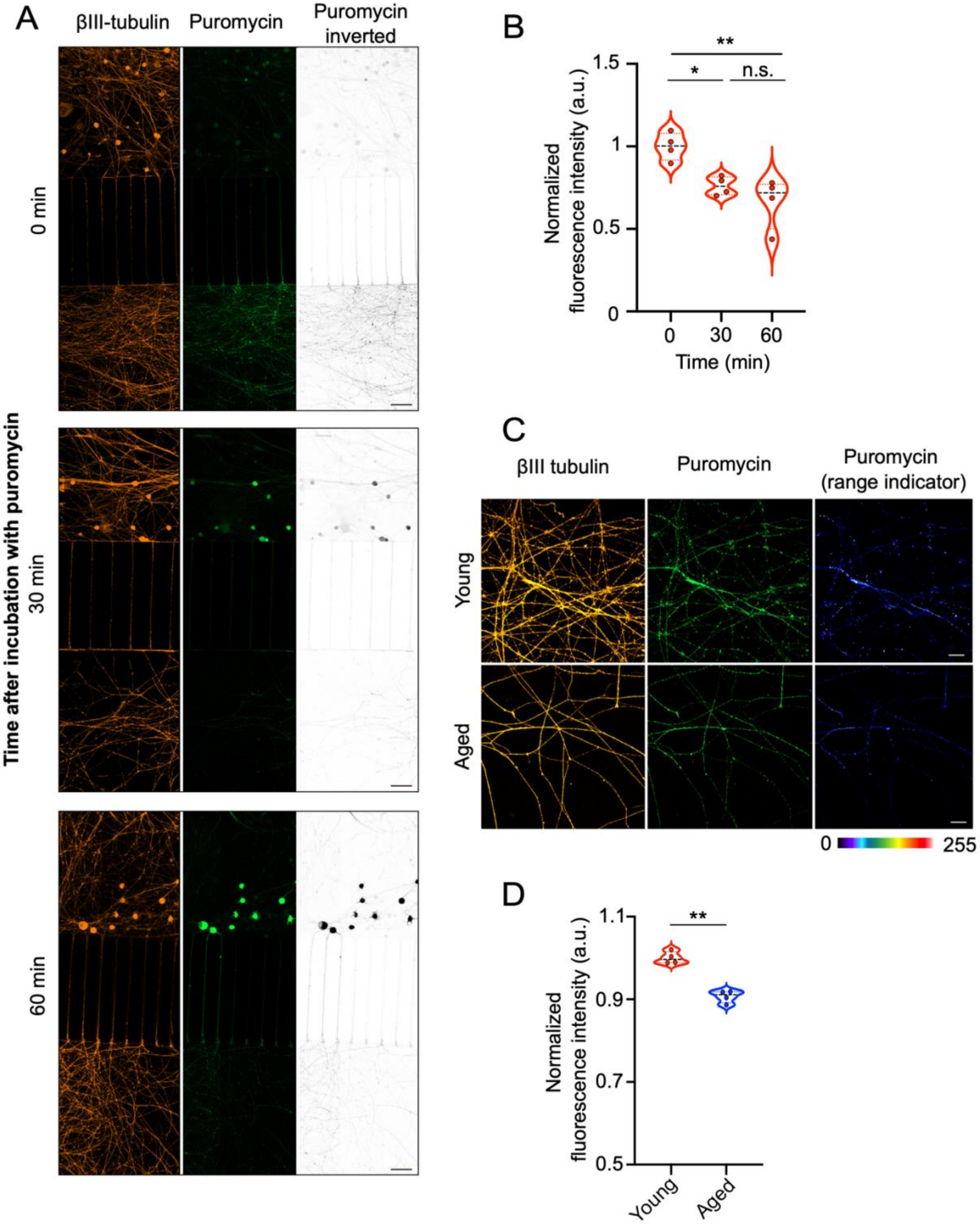

### Supplementary Figure 4

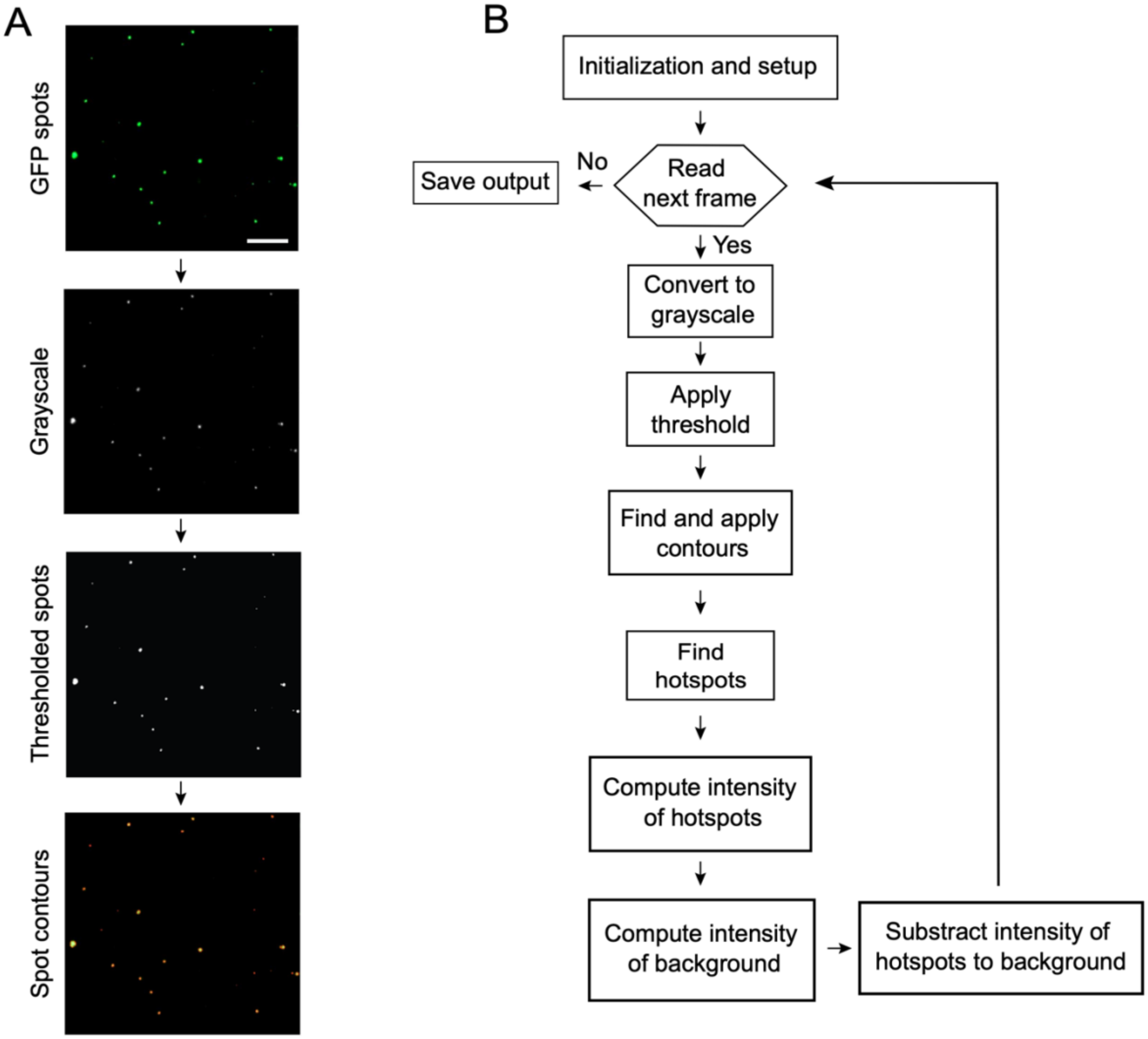

### Supplementary Figure 5

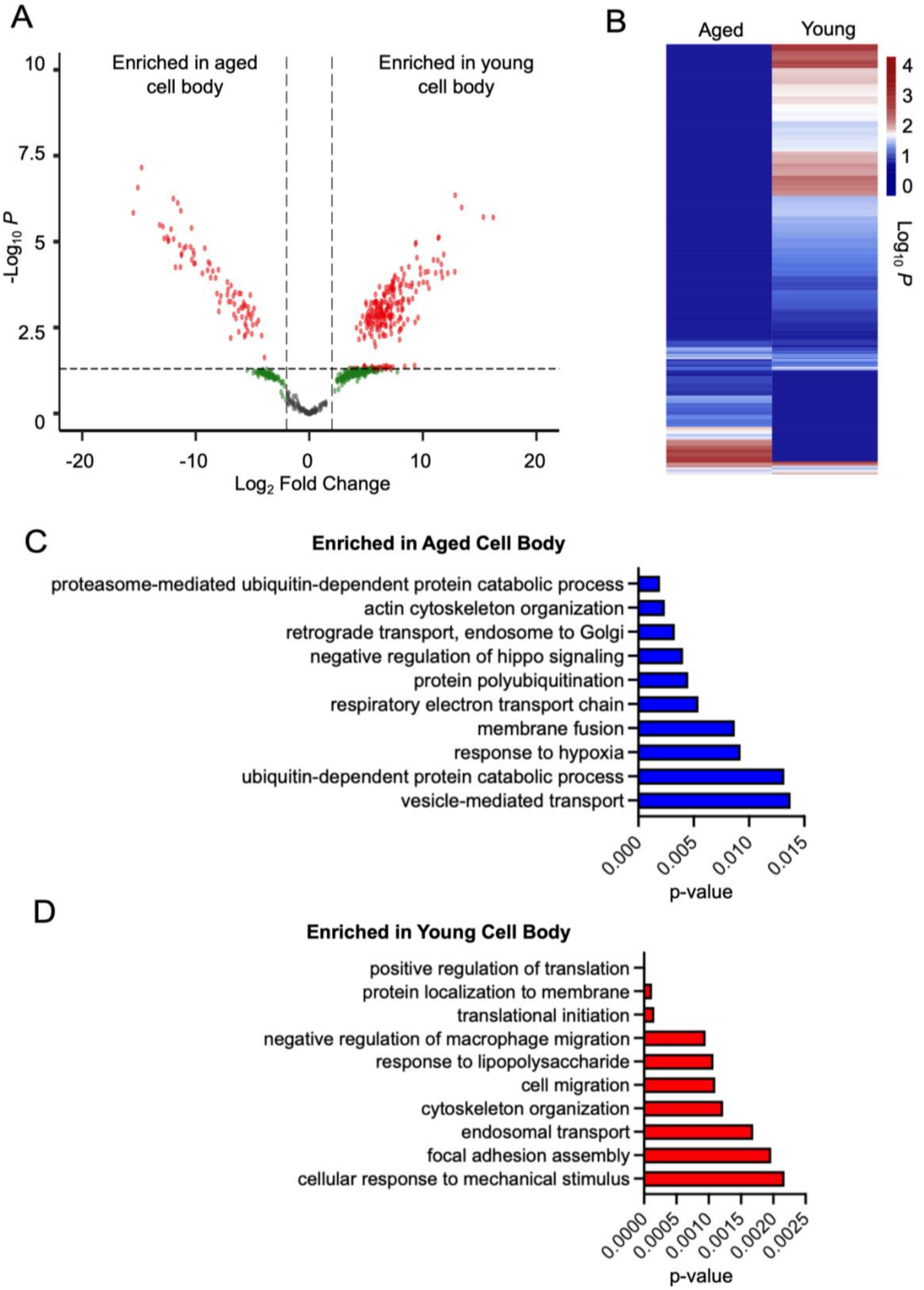

### Supplementary Figure 6

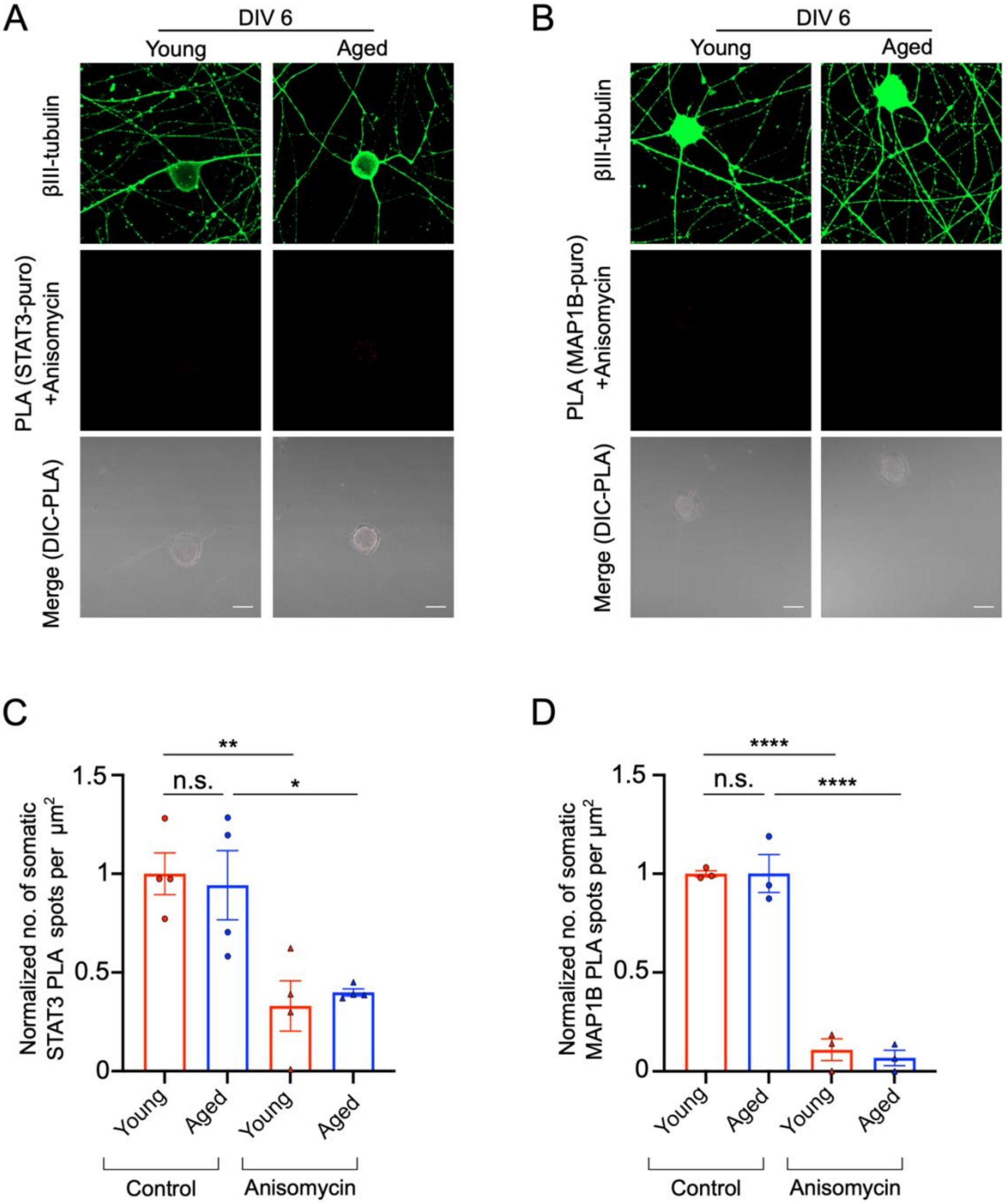

### Supplementary Figure 7

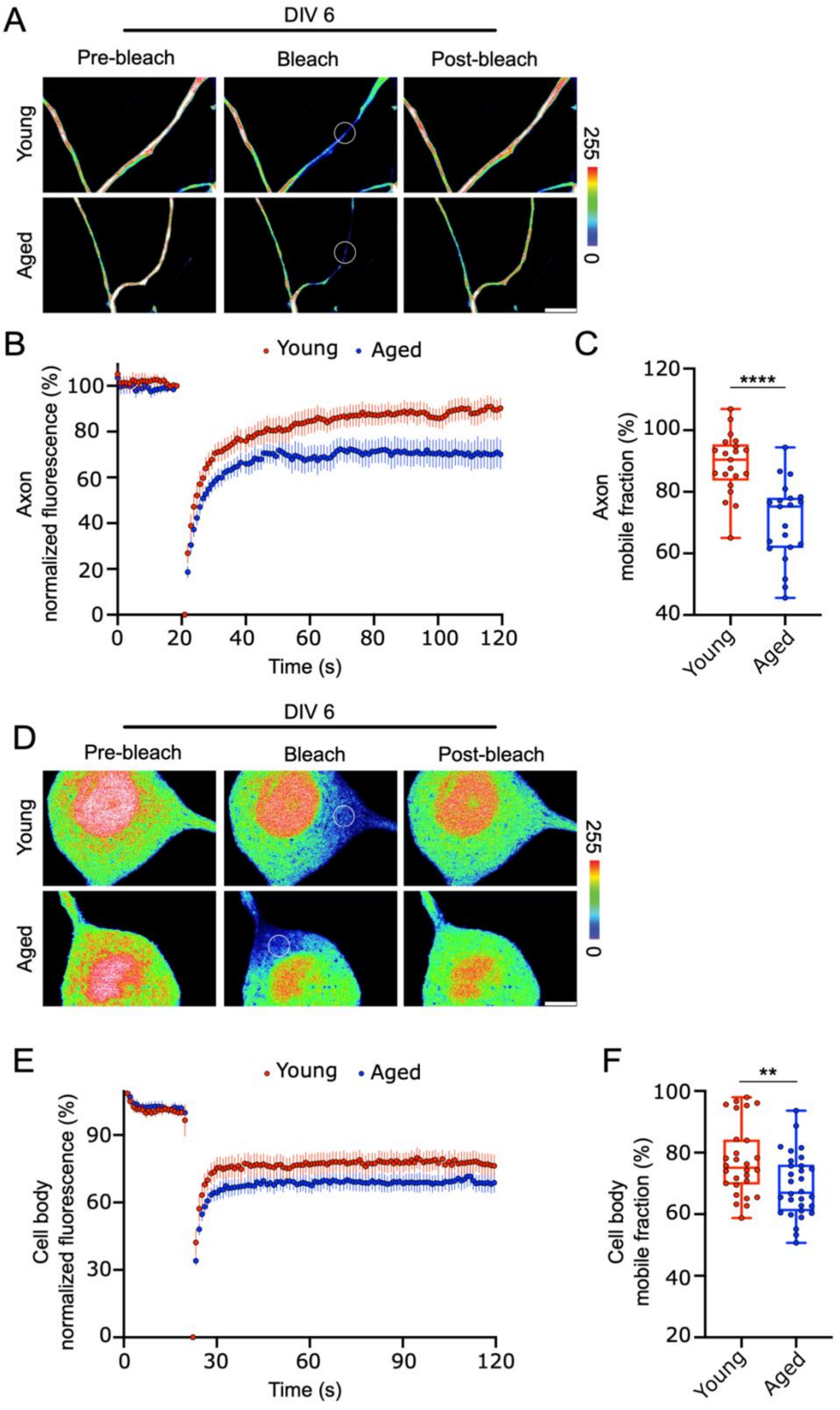

### Supplementary Figure 8

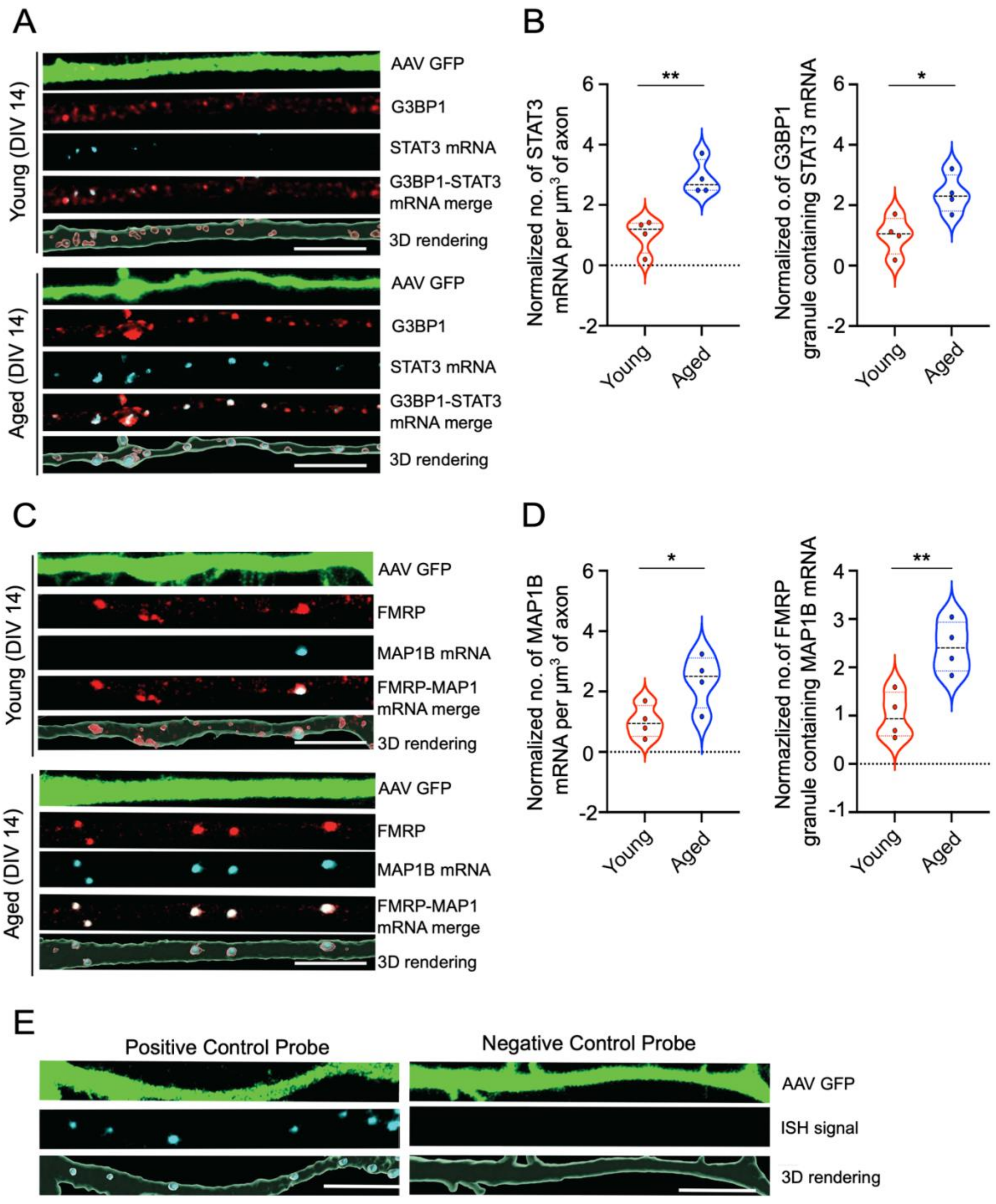

#### Supplementary Figure 9

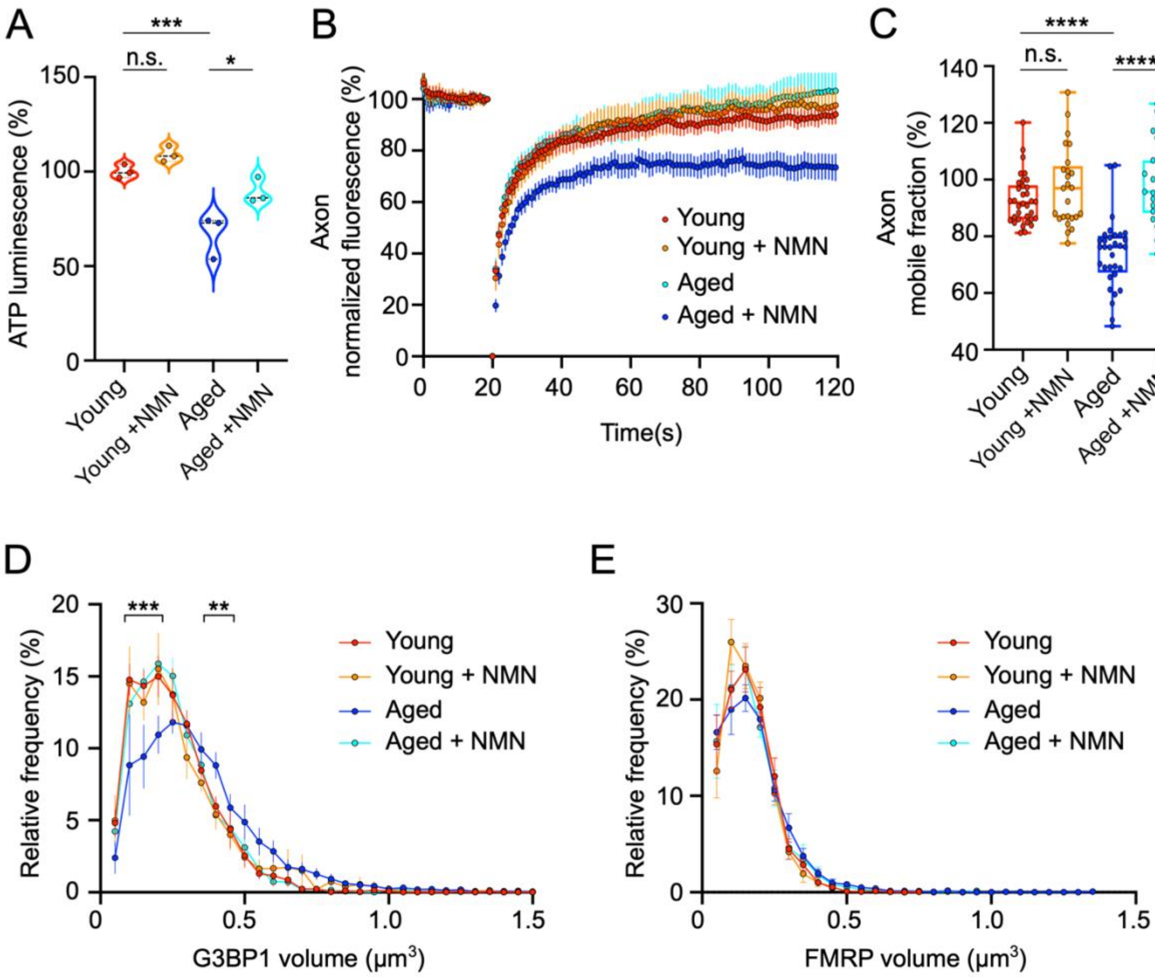

### Supplementary Figure 10

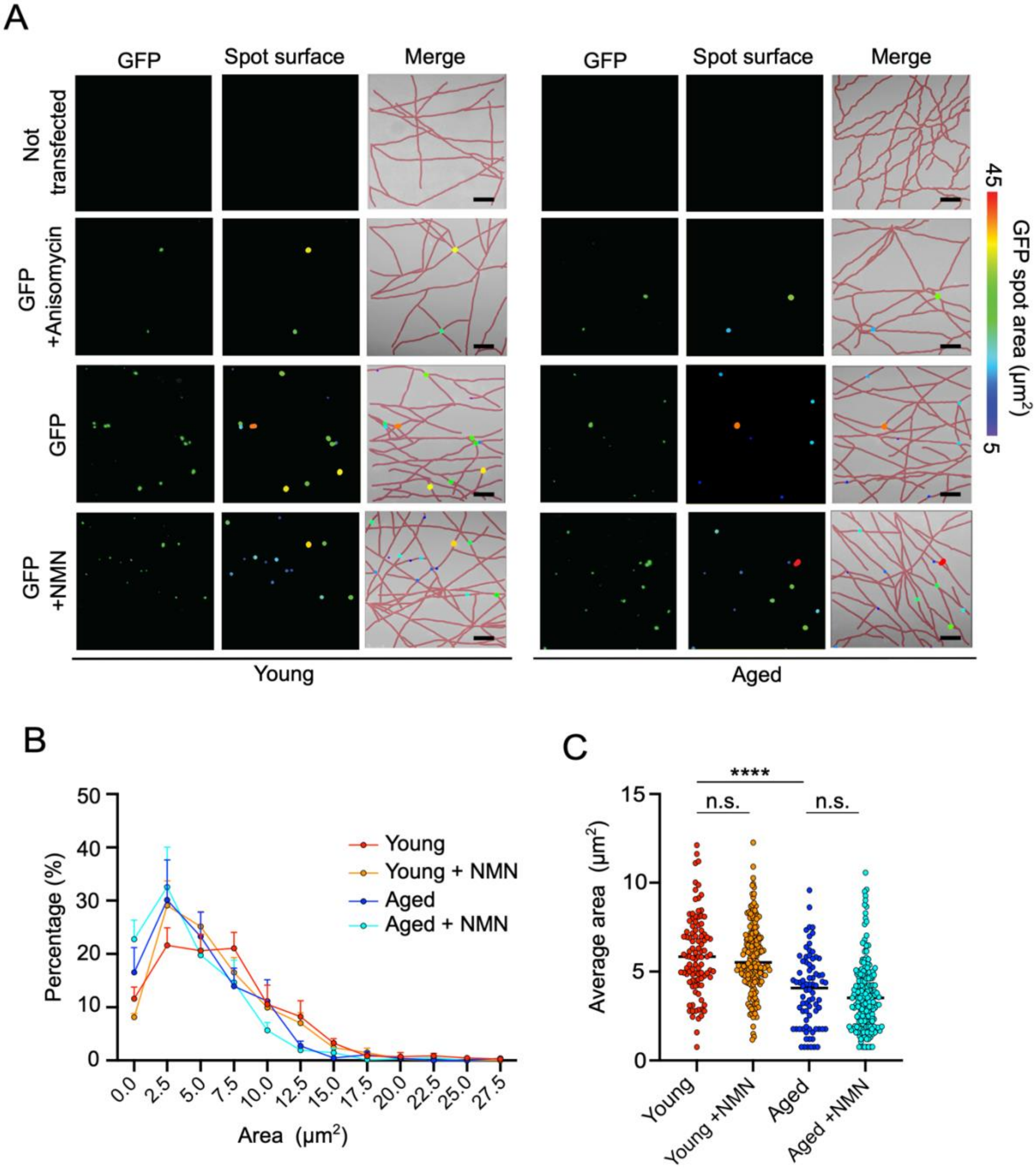

#### Supplementary Figure 11

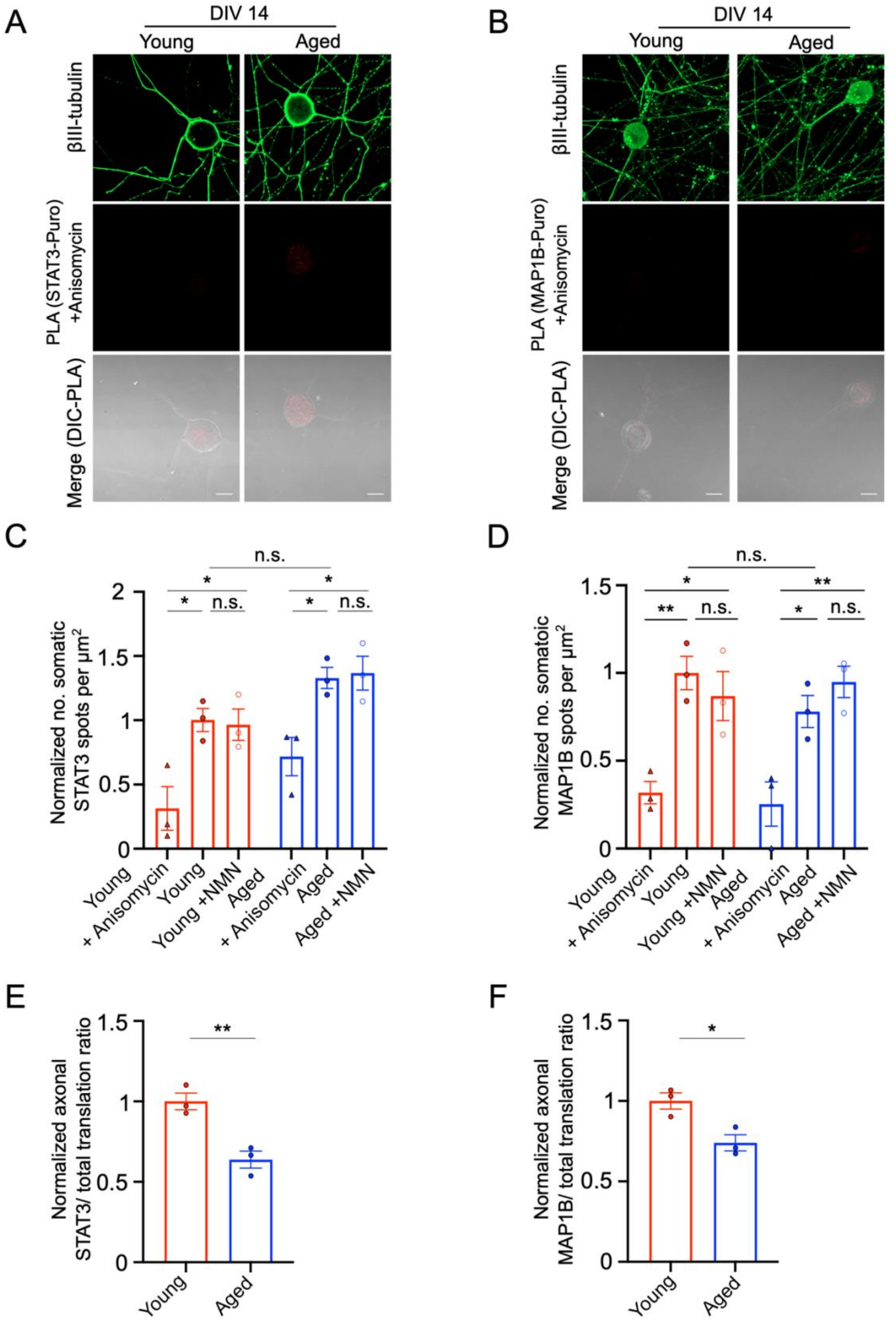
